# EVQuant; high-throughput quantification and characterization of extracellular vesicle (sub)populations

**DOI:** 10.1101/2020.10.21.348375

**Authors:** T.A. Hartjes, J.A. Slotman, M.S. Vredenbregt, N. Dits, R. Van der Meel, D. Duijvesz, J.A. Kulkarni, P.J. French, W.A. Van Cappellen, R.M. Schiffelers, A.B. Houtsmuller, G.W. Jenster, M.E. Van Royen

## Abstract

Extracellular vesicles (EVs) reflect the cell of origin in terms of nucleic acids and protein content. They are found in biofluids and represent an ideal liquid biopsy biomarker source for many diseases. Unfortunately, clinical implementation is limited by available technologies for EV analysis. We have developed a simple, robust and sensitive microscopy-based high-throughput assay (EVQuant) to overcome these limitations and allow widespread use in the EV community. The EVQuant assay can detect individual immobilized EVs as small as 35 nm and determine their concentration in biofluids without extensive EV isolation or purification procedures. It can also identify specific EV subpopulations based on combinations of biomarkers and is used here to identify prostate-derived urinary EVs as CD9-/CD63+. Moreover, characterization of individual EVs allows analysis of their size distribution. The ability to identify, quantify and characterize EV (sub-)populations in high-throughput substantially extents the applicability of the EVQuant assay over most current EV quantification assays.

## Introduction

Extracellular vesicles (EVs) are a large family of bilipid membrane vesicles that are secreted by most if not all cells and can be found in many biological fluids^1, 2^. Due to their biogenesis, EVs not only contain membrane bound proteins but also a fraction of the cytoplasm that represents the (malignant) cells from which they originate. Disease-related RNAs and proteins present in secreted EVs’ lumen or on their surface can serve as powerful indicators in body fluids for disease elsewhere. As these ‘liquid biopsies’ are easily obtained without the need for invasive biopsies or surgery, EVs represent an ideal source for diagnostic, prognostic and monitoring biomarkers in a whole range of diseases including cancer^3–7^. However, detection and quantification of EVs in biofluids is challenging because of their size and heterogeneity, but also because lipoproteins and protein complexes have the same size range and have an overlap in composition and morphology^8–15^. Most of current approaches for EV quantification are laborious measurements and require expensive, highly specific, sometimes custom-made equipment. Moreover, EV quantification by most techniques are biased towards certain size or concentration ranges and/or requires EV isolation or purification leading to EV loss and underestimation of EV concentration. As a result, many of the current approaches are less suitable to quantify and characterize the full spectrum of EVs, reflected by the large variations found in literature ^16, 17^. More importantly, most technologies are not able to detect and quantify specific EV subpopulations based on biomarkers present on individual vesicles. Clearly, there is still a need for improved methods to efficiently quantify and characterize EVs, especially for moving toward a clinical setting.

Our aim was to develop a rapid, easy to use, widely accessible, and cost-effective EV quantification and characterization approach. We describe a novel microscopy-based assay, which allows multicolor fluorescence-based detection of individual EVs to quantify total (EVs/ml) and biomarker-specific EV populations present in several biological fluids, without the need for extensive EV isolation and purification procedures. This assay opens up a range of possibilities for EV biomarker analysis in both research and clinical settings.

## Materials and Methods

### Cell culture and media collection

All cell lines used in this study were cultured using standard culture conditions (Supplemental Table S5). For direct EV quantification in conditioned media or EV isolates, cells were cultured in a T175 (175 cm^2^) culture flasks (Greiner Bio-One, Frickenhause, Germany) in complete media. When cells reached 80% confluency, cells were washed with PBS (Lonza, Verviers, Belgium) and grown in corresponding FCS-free medium for 48 hours to avoid FCS-derived EV contamination in the analysis. Cell-conditioned media was collected, centrifuged at 3,000xg for 20 minutes at 4°C and either used directly for EV quantification and/or EV isolation or stored at −80°C for long term storage.

### Urine collection and storage

Whole urine from men with or without prostate cancer was collected at the out patients clinic from the Erasmus Medical Center Rotterdam (Medical Ethics Approval number 2005-077 and 2010-176). Urines from men were collected after digital rectal examination (DRE) or without DRE. Furthermore, whole urines were also collected from men after radical prostatectomy and female volunteers. All urines were temporarily stored at 4°C after collection and processed within 1 day. Urine was centrifuged at 3,000xg for 20 minutes at 4°C to remove dead cells and cellular debris and stored at −80°C for long term storage.

### Extracellular vesicle isolation

EVs were isolated using ultra-centrifugation according to the previously described protocol by Duijvesz et al.^39^. Cell conditioned media was subjected to consecutive centrifugation steps. First, cells and cell debris were removed by centrifugation for 20 min at 3,000xg at 4°C. Next, larger vesicles were depleted by centrifugation for 30 minutes at 10,000xg at 4°C (SW28 rotor, Beckman Coulter). In the final step, EVs were pelleted by ultracentrifugation at 100,000xg for 120 minutes at 4°C (SW28 rotor, Beckman Coulter). Additionally, the EV pellets were resuspended in 3 ml 0.32 M sucrose (diluted in PBS) solution and centrifuged again at 100,000xg for 60 minutes at 4°C (SW60 rotor, Beckman Coulter). Sucrose supernatant was removed and pellet was resuspended in 500 μl phosphate-buffered saline containing 0.01% bovine serum albumin (0.2 μm filtered) and stored at −80°C.

### Preparation of liposomes

Liposomes were prepared using rapid-mixing methods as described before by Kulkarni *et al.*^40^. 1,2-distearoyl-sn-glycero-3-phosphocholine (DSPC, Avanti Polar Lipids), cholesterol (Sigma) and 1,2-distearoyl-sn-glycero-3-phosphoethanolamine-N-[methoxy(polyethylene glycol)-2000] (PEG-DSPE, Avanti Polar Lipids) were dissolved in ethanol in a molar ratio of 53:42:5 to a final concentration of 10 mM total lipid (Batch 1; Supplemental Table S2). The lipids in ethanol solution and 25 mM acetate buffer pH 4.0 were mixed using microfluidic mixers (Precision Nanosystems) at flow rate ratios of 1:1 and 1:3 respectively with a combined final flow rate of 12 mL/min. For the second batch (Batch 2; Supplemental Table S4), organic and aqueous solutions were mixed using a T-junction mixer at flow ratios of 1:1, 1:2 and 1:3 were used. Formulations were dialyzed against PBS pH 7.4 overnight. LNPs were passed through 0.2 μm filters, concentrated by 12-14 kDa cutoff Vivaspin centrifugal concentrators and stored at 4°C. Size distribution was determined by dynamic light scattering and zèta potential by electrophoretic light scattering using a Zetasizer Nano ZS (Malvern). Lipid concentration was determined by Cholesterol E assay (Wako).

### EVQuant assay

Nanoparticles are provided and/or diluted in a standard end volume (40 μl) and if applicable fluorescently labeled. EVs are either non-specifically labeled using the fluorescent membrane dye Octadecyl Rhodamine B Chloride R18 (Life Technologies, catalog# O246, final staining concentration of 0.33 ng/ul) or PKH26/67 dye (Sigma-aldrich, catalog# MINI26-1KT/MINI67-1KT, final staining concentration of 5 μM) and/or specifically labeled using the immunofluorescent antibodies Mouse CD9 Monoclonal (MEM-61)-Alexa Fluor® 647 antibody (Thermo Fisher, Catalog# MA5-18154, 1:25 dilution, 50 μl staining volume) and/or Mouse CD63 (MX-49.129.5)-Alexa Fluor®488 antibody (Santa Cruz, catalog# sc-5275 AF488, antibody concentration 2.5 μg/ml, 50 μl staining volume). Subsequently, without any isolation or purification procedures, nanoparticles are immobilized in a non-denaturing polyacrylamide gel (final ratio of 16% (w/w) acrylamide/bis-acrylamide) to allow detection of low intensity signals. Images are acquired by confocal microscopy to minimize auto-fluorescence of unlabeled dye/antibody, using a conventional laser scanning confocal microscope (CLSM510Meta, Zeiss) or a high-content screening system (Opera Phenix, Perkin Elmer). Instrument specifications are found in Supplemental Information. Phosphate buffered saline (PBS) or serum free culture medium, supplemented with the same concentration of fluorescent dye/antibody are used as negative controls. Cell-conditioned medium from DU145 prostate cancer cells from which aliquots are stored at −80°C and 100 nm liposomes or 100 nm fluorescent tetraspeck beads (Thermo Fischer, catalog# T7279) are used as internal standards for quality control. Multiple images are acquired for each sample and fluorescent signals are quantified using the in house developed EVQuant Plugin (latest version will be available at the open source repository https://github.com/MEvanRoyen/EVQuant) for the open source software Fiji or the Opera Phenix analysis software (Harmony 5.4, Perkin Elmer)^41^. To quantify the number of particles in the images, particles are selected and counted based on the non-specific membrane labeling. Particles are counted using the “Find Maxima” (Fiji) or “Find Spots” (Harmony Software) function which determines local maxima in an image using a threshold. Absolute quantification of the particles requires a one-time calibration of the detection volume used for quantification on each specific confocal microscope which is explained below. When detection volume is known, absolute particle concentration is calculated by the formula presented below. A detailed protocol for EVQuant analysis is found in Supplemental Information.

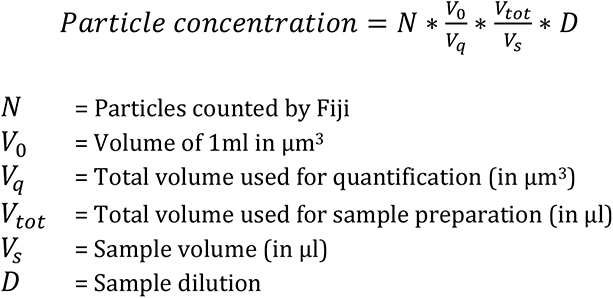

### Calibration of detection volume used by EVQuant

Because of its three-dimensional context, absolute quantification by the EVQuant assay requires a one-time calibration of the detection volume in which spots are detected for each microscope, as it depends on the unique optical specifications and settings of the (confocal) microscope that is used. Due to the scattering of light, resulting in the typical point spread function (PSF), EVs will be detected in multiple planes (Supplemental Figure S1A). To analyze the effect of the PSF on the effective optical slice thickness (EOST) in single plane measurements, multiple Z-stacks need to be acquired to quantify both detected particles per plane and particle centers per plane. Samples of interest (e.g. Beads, EVs) are prepared using the standard EVQuant protocol and multiple 10-20 μm z-stacks (interval of 0.5 μm) are acquired and analyzed using the EOST calibration option in the EVQuant plugin. First, particles in each plane are selected and quantified based on the non-specific membrane labeling. Next, analysis of individual particles in the stack of images is used to quantify the number of particles in 3D. Using the 3D defined particles, we can determine the actual number of detected particles and assign a center for each particle to the corresponding plane of the z-stack. Finally, the effective optical slice thickness (EOST) is calculated by dividing the average number of detected particles (2D) in each plane by the average detected particle centers (3D) in each plane and multiplying it by the slice interval which was used in the z-stack (0.5 μm). The known EOST (and thus the detection volume) of the microscope system used, now allows absolute particle quantification. A detailed protocol for the calibration of the detection volume is found in Supplemental Information.

### Quantification of EV subpopulations using immunofluorescent antibodies in cell line derived EVs and urine

To quantify EV subpopulations based on combinations of specific biomarkers, samples including negative controls, are prepared using the standard EVQuant protocol. For each sample, multiple images (multichannel) are acquired and fluorescent signals are analyzed using the EVQuant plugin. Expression of a specific protein on the EV surface is analyzed as follows. First, particles are selected based on the non-specific membrane labeling. Next, mean fluorescent intensity of each marker of interest in every detected spot is calculated. Fluorescent intensity of each marker in the area surrounding each spot is also calculated and subtracted from the mean intensity in the spot (relative intensity). For each biomarker, the values of spots that had a negative relative intensity are selected and combined with absolute values of the same negative relative intensities to create a gaussian distribution of background noise for each sample. Distribution of background noise for each biomarker is plotted in a histogram and fitted using a Gaussian function. Mean plus three times the standard deviation is used as a threshold to define EVs being positive for each marker of interest. The number of EVs having each specific biomarker combination is determined and corrected for the number of particles present in the Negative Control (NC) sample. Fraction of each specific subpopulation is calculated by dividing the number of particles positive for the marker combination to the total number of detected particles. A detailed protocol for EVQuant analysis using the combination of non-specific and immune-specific labeling is provided in Supplemental Information.

### Quantification of EV subpopulations in serum and plasma using immunofluorescent antibodies

Because the non-specific membrane labeling cannot be used to select EVs due to the high background signals from lipoproteins present in serum and plasma, particles are selected based on the fluorescent signals of the biomarkers themselves. To quantify EV subpopulations in serum and plasma based on combinations of specific biomarkers, samples including negative controls, are prepared, imaged using the standard EVQuant protocol using antibodies and analyzed by Fiji. For each biomarker channel, particles are selected using the find maxima function, followed by creating a binary mask of the detected particles for this channel. The presence of combinations of biomarkers is then determined by combining the masks of the different biomarkers and analysis of the overlapping signals. The number of EVs having each specific biomarker combination is determined and corrected for the number of particles present in the Negative Control (NC) sample.

### EV size characterization

Similar to EV quantification, EV size characterization requires a one-time calibration for each microscope, as it depends on the unique optical specifications and settings of the (confocal) microscope that is used. Size analysis is calibrated using a liposome calibration sample with a known particle size (i.e. determined by DLS/EM). To determine the particle size distribution, samples including the calibration sample and negative control, are prepared using the standard EVQuant protocol and multiple 10-20 μm z-stacks (interval of 0.5 μm) per sample are acquired. First, the stacks of the calibration sample are analyzed using the sizing calibration option within the EVQuant plugin followed by size analysis of all samples (including the calibration sample). In short, the maximum intensity of every individual particle is determined in 3D and therefore independent of the focal plane. To convert fluorescent intensity to particle size, the following analysis is only performed in the calibration plugin but not in the sizing plugin. The maximum intensities in Z of all particles are plotted in a histogram and fitted using a gaussian to determine the mean fluorescent intensity of the calibration particles. Theoretically, fluorescent intensity of the particle increases quadratic with size. The size calibration factor is therefore calculated by dividing the mean fluorescent intensity through the squared radius of the calibration particles. This calibration factor is used in the sizing plugin to determine the size of every detected particle in the rest of the samples. A detailed step-wise protocol for the EV size characterization is found in Supplemental Information.

### Nanoparticle tracking analysis (NTA)

EV quantification and characterization was performed by nanoparticle tracking analysis (NTA) using the NanoSight NS3000 system (Malvern, Sysmex) in light scatter mode using the 405 nm laser. Fluorescence mode could not be used on our unpurified samples due to background fluorescence from unbound dye. Samples were thawed, diluted in 0.2 μm filtered PBS to instrument specific concentrations (50-100 particles in the field of view) and three consecutive measurements were performed for 60 seconds at controlled temperature of 25°C. Previous studies in which different EV quantification techniques were extensively compared showed NTA yielded the highest EV concentrations but also showed that measured EV concentrations are highly dependent on instrument settings^16^. Therefore, we evaluated the effect of camera level and detection threshold on the quantification of EVs on our Nanosight NS300 (concentration update installed) by acquiring movies at a range of camera levels (10 to 14 at preprogrammed NTA settings) and processing the movies using different detections thresholds (2 to 8). Only small differences in EV concentrations were observed between camera level 10 to 13, even when different detection thresholds were used. However, differences became bigger when using camera level 14 (Supplemental Figure S1). When using camera level 14, images started to become too bright for accurate analysis. As a higher camera level would be most favorable for detecting small vesicles, all measurements were performed using camera level 13 and processed using detection threshold 3 in the NTA software.

### TR-FIA

EV biomarker analysis using a human specific time-resolved florescent immunoassay was performed as previously described by Duijvesz et al.^42^. In short, a streptavidin-coated 96-wells plate (KaiSA96, Kaivogen, Turku, Finland) was incubated with either human-specific biotinylated CD9 or CD63 antibody (200 ng / 100 μl in each well) for 1 hour while shaking (750 rpm) at RT. The unbound antibodies were removed by washing three times using a washing buffer (catalog# 42-02, Kaivogen, Turku, Finland) and an automated plate washer (TECAN Columbus). EV containing samples were diluted in a sample buffer and 100 ul volume was added to each well of the 96-wells plate (samples were measured in triplo). Samples were incubated for 1 hour while shaking at RT which allowed the EVs being captured on the surface by CD9 or CD63. Unbound EVs were removed and plate was washed three times. For EV detection, a second antibody which was labeled with europium was added to the plate (25 ng Eu-labeled antibody / 100 ul in each well) and incubated for 1 hour at RT. Unbound antibodies were removed by washing three times. At last, 100 μl of enhancement solution (Perkin Elmer) was added to each well and incubated for 15 minutes while shaking at RT. Europium counts (time-resolved fluorescence) was measured using a Wallac Victor 2, 1420 mulilabel counter (Perkin Elmer). In this study only two combinations of capture and detection antibodies were used; CD9-CD9 and CD63-CD63, respectively.

### Cryogenic transmission electron microscopy (cryo-TEM)

Cryo-TEM was performed as described previously^43^. Briefly, 2-5 uL of concentrated LNP (20-25 mg/mL total lipid) was added to Lacey-formvar copper grids, and plunge-frozen using a FEI Mark IV Vitrobot (FEI, Hillsboro, OR). Grids were stored in liquid nitrogen until imaged. Grids were moved into a Gatan transfer station pre-equilibrated to at least −180°C prior to add grids to the cryogenic grid holder. A FEI LaB6 G2 TEM (FEI, Hillsboro, OR) operating at 200 kV under low-dose conditions was used to image all samples. A bottom-mount FEI Eagle 4K CCD camera was used to capture all images at a 47-55,000x magnification with a nominal under-focus of 1-2 μm to enhance contrast. Sample preparation and imaging was performed at the UBC Bioimaging Facility (Vancouver, BC).

## Results

### Detection and quantification of individual membrane vesicles by EVQuant

The EVQuant assay counts individual fluorescently-labeled (Rhodamine-R18) vesicles immobilized in a transparent matrix by confocal microscopy without the need to remove free dye. The transparent matrix avoids detection of reflected excitation light, limits background fluorescence of dye bound to the glass surface and allows long exposure times for sensitive vesicle detection (Figure 1a). EVQuant enables detecting a variety of synthetic vesicles like liposomes and polymersomes, but also biological EVs in minimally processed cell conditioned medium and body fluids (i.e. after centrifugation at 3,000xg for 20 minutes at 4°C) (Figure 1b). As expected, the dye-only control only showed a limited number of false-positive spots (Figure 1c). To be able to express the EV detections as an absolute EV concentration, we calibrated the imaging volume in which EVs are detected. The microscope system specific (here Zeiss CLSM confocal microscope, details are described in the materials and methods section) effective optical slice thickness (EOST) is 1.67 μm (Supplemental Table S1). The EOST calibration is validated using a 100 nm tetraspeck polystyrene bead sample with a known concentration provided by the manufacturer (Invitrogen) (Supplemental Table S1, Supplemental Figure S1a-b).

**Figure 1.**
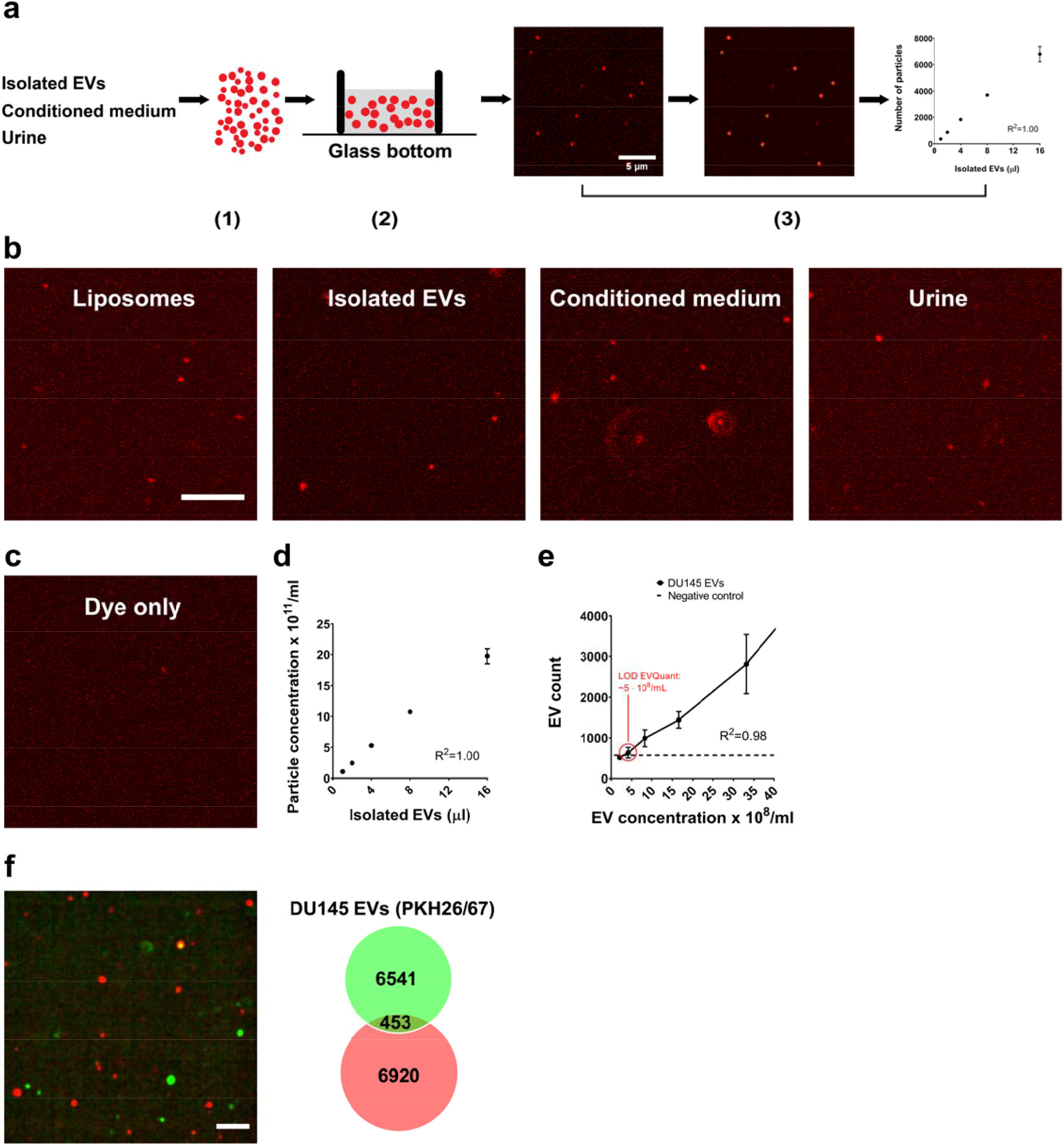
Detection and quantification of individual membrane vesicles. **(a)** Workflow of EVQuant: Membrane particles, for example in EV isolates or minimally processed conditioned medium or urine, are fluorescently labeled by short incubation with a fluorescent dye and/or primary-labeled immunofluorescent antibodies (1). Subsequently, without removal of free fluorescent dyes, the membrane particles are immobilized in a standard non-denaturing polyacrylamide gel inside a sample holder that is compatible with confocal microscopy (2). In the last step, images are acquired using fluorescent confocal microscopy and analyzed using the open source software Fiji and the EVQuant plugin for absolute particle quantification (3). **(b)** Representative images of non-specific fluorescently labeled individual synthetic and biological vesicles detected by EVQuant on a LSM510 laser scanning confocal microscope (Zeiss). Scale bar 5 μm. **(c)** Representative image of the dye only control (PBS). **(d)** Absolute EV concentrations of serial two-fold dilutions of isolated EVs from DU145 prostate cancer cells measured by EVQuant. **(e)** Limit of quantification (LOD) of the EVQuant assay was calculated using serial two-fold dilutions of isolated EVs from DU145 prostate cancer cells. Dotted line represents the average number of detected particles (counts) in a series of negative controls. **(f)** DU145 EVs were split in two and either labeled with the fluorescent membrane dye PKH26 (red) or PKH67 (green). After fluorescent labeling and removal of free dye, EVs were pooled and measured by EVQuant (left). Venn diagram showing the number of detected PKH26, PKH67 or double positive EVs in the acquired images (right). Scale bar 10 μm. EVQuant is performed in duplicate (Mean +- SD)

Quantification of isolated EVs from a prostate cancer (PCa) cell line (DU145), showed that measured EV concentrations were linearly related to dilution suggesting the detection of individual EVs and showed that the lowest concentration of EVs that can be detected (limit of detection or LOD) is ~5E8 particles/ml (Figure 1d,e). To verify the detection of individual EVs, isolated EVs from cell-conditioned media were labeled by either red or green fluorescent membrane dyes (PKH26 or PKH67), mixed and analyzed by EVQuant. The detected EVs were either PKH26 or PKH67 positive with a minimal percentage (ca. 3%) of two-color positive detections, confirming that EVQuant is able to detect individual EVs (Figure 1f).

The EVQuant assay is even able to efficiently detect relatively homogeneous populations of 40 nm synthetic liposomes as determined by dynamic light scattering (DLS) and Cryo-electron microscopy (Cryo-EM) (Figure 2a-b; Supplemental Figure S2a-b, Supplemental Table S2). Interestingly, EVQuant analysis showed a 2.6-fold higher concentration of LIPO1 (80 nm) and even a 5.6-fold higher concentration of LIPO2 (40 nm) compared to NTA (NanoSight NS300, Sysmex) (Figure 2c). With that, the EVQuant (but not the NTA) measurements correlate well with the higher liposome concentration in LIPO2 compared to LIPO1 sample, based on the higher lipid concentration together with the obvious smaller number of lipids in individual smaller liposomes (Supplemental Table S2). This observation and the significantly larger size of the LIPO2 liposomes determined by NTA compared to DLS and Cryo-EM, strongly suggests that NTA is less sensitive in detecting (smaller) EVs than DLS, Cryo-EM and also EVQuant (Supplemental Figure S2a).

**Figure 2.**
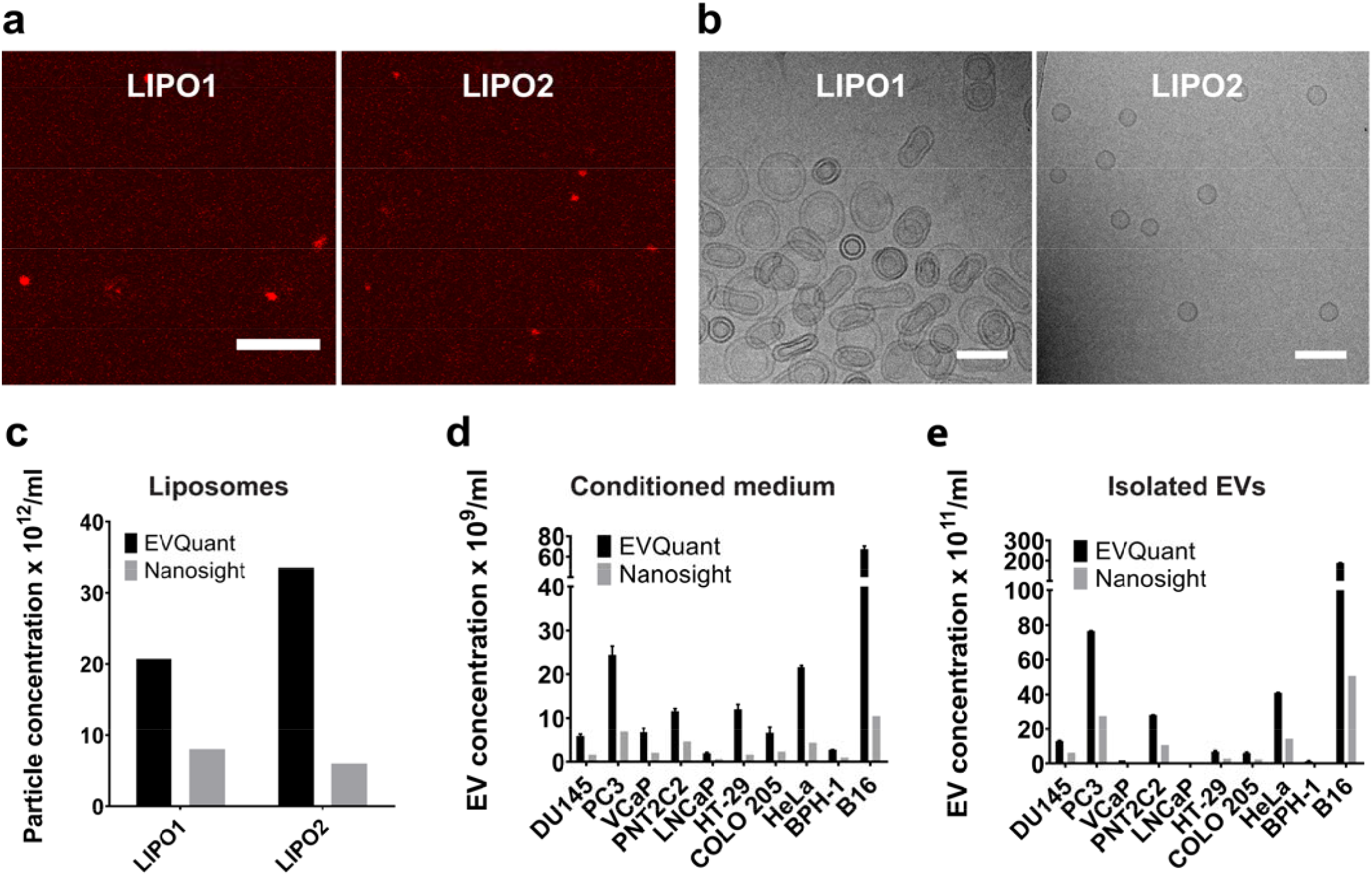
Quantification of membrane particles by EVQuant and NTA (NanoSight NS300). **(a)** Representative images of lipid nanoparticles of around 80 and 40 nm in size (LIPO1 and LIPO2 respectively) acquired by Cryo-TEM. Scale bar 100 nm. **(b)** Representative images of detected fluorescent lipid nanoparticles LNP1 and LNP2 acquired by EVQuant using a laser scanning confocal microscope (LSM510, Zeiss). Scale bar 5 μm **(c)** Comparison between quantification of LIPO1 and LIPO2 lipid nanoparticles by EVQuant and NTA. **(d-e)** EV concentration in minimally processed cell conditioned medium and matching EV isolates of a panel of 10 cell lines from different origins was quantified by EVQuant and NTA. Single experiment was performed for NTA and EVQuant was performed in duplicate (Mean +- SD).

Together, these data demonstrate that EVQuant is able to efficiently detect individual EVs as small as 40 nm and accurately quantifies EVs over a large concentration range. Importantly, EV detection was not hampered by background fluorescence of free dye, making time-consuming EV isolation/purification procedures unnecessary.

### Quantification of EVs in biological samples

To avoid underestimation and/or bias, EVs in complex biological samples are preferably quantified without performing extensive isolation and purification procedures. Similarly to isolated EVs, serial dilutions of minimally processed cell-conditioned medium from VCaP cells showed a linear relation between measured concentrations and dilution (Supplemental Figure S3a). EV quantification in minimally processed cell conditioned media and matching EV isolates from a panel of cell lines (n=10) showed a strong correlation between EVQuant and NTA analyses (Supplemental Figure S3b-c). However, concentrations determined by EVQuant in general were on average 3.4-fold (2.1-7.3) higher, again indicating a higher sensitivity of EVQuant (Figure 2d-e). Importantly, only up to 17% of the vesicles were recovered by EV isolation using a standard ultracentrifugation protocol and together with a high variety in recovery rate, suggests that minimally processed samples are strongly preferred over EV isolates for EV quantification (Supplemental Figure S3d).

To enable (semi-)high-throughput EV quantification, we also implemented EVQuant on a confocal microscope-based high-content screening system (Opera Phenix, Perkin Elmer) with a calibrated EOST of 2.17 μm (Supplemental Table S3). Measurement of the same samples on both microscope systems resulted in very similar EV concentrations, demonstrating that EVQuant can easily be implemented on a variety of confocal microscopy-based systems including (semi-)high-throughput systems (Supplemental Figure S3e).

### Quantification of EV subpopulations

The use of EVs for biomarker analysis will rely on detecting (combinations of) specific proteins on the EV surface. Quantification of human CD9 and CD63 EV surface markers in EV isolates from 9 cell lines (mouse B16 as negative control) using the highly sensitive time-resolved fluorescence immunoassay (TR-FIA)^12^ showed high variation between cell lines (Figure 3a; Supplemental Figure S4a). This could be explained by variation in EV concentration but also by variation in the presence of biomarker molecules on individual EVs. Obviously, bulk analyses (like TR-FIA) do not discriminate between these options, however EV concentrations determined by EVQuant showed no strong correlation with the marker profiles, suggesting variation in biomarker presence among the cell lines (Figure 3b, Supplemental Figure S4a-c). To investigate this further, we combined the general membrane labelling with specific immune-detection of the tetraspanin CD9 and CD63 surface markers in a multicolor EVQuant analysis to determine marker positive subpopulations (Figure 3c). Total EV quantification based on the non-specific membrane dye was not affected by additional labelling with immunofluorescent antibodies (Supplemental Figure S4d). Overall, a larger fraction of EVs is positive for CD9 compared to CD63, which is in line with TR-FIA analysis (Figure 3a,d). However, no strong correlation was found between concentration of CD9 and CD63 sub-populations and corresponding europium counts by TR-FIA (Supplemental Figure S4e-f). Again, this could well be explained by the difference in signal detection by the two assays: more epitopes on the surface of an EV will increase the TR-FIA signal, but more epitopes on detected EVs in the EVQuant assay will not result in a higher EV count. To compare the relative number of epitopes between the cell-lines using EVQuant, we determined the median CD9 and CD63 fluorescent intensities from individual EVs (Supplemental Figure S4g). For CD9, the median spot intensity correlated well (R^2^=0.87) to the normalized europium counts suggesting that there are indeed differences in numbers of epitopes on EVs between the cell lines (Supplemental Figure S4h). Due to very low number of CD63-positive EVs in some of the samples, no correlation was found for CD63 (Supplemental Figure S4i).

**Figure 3.**
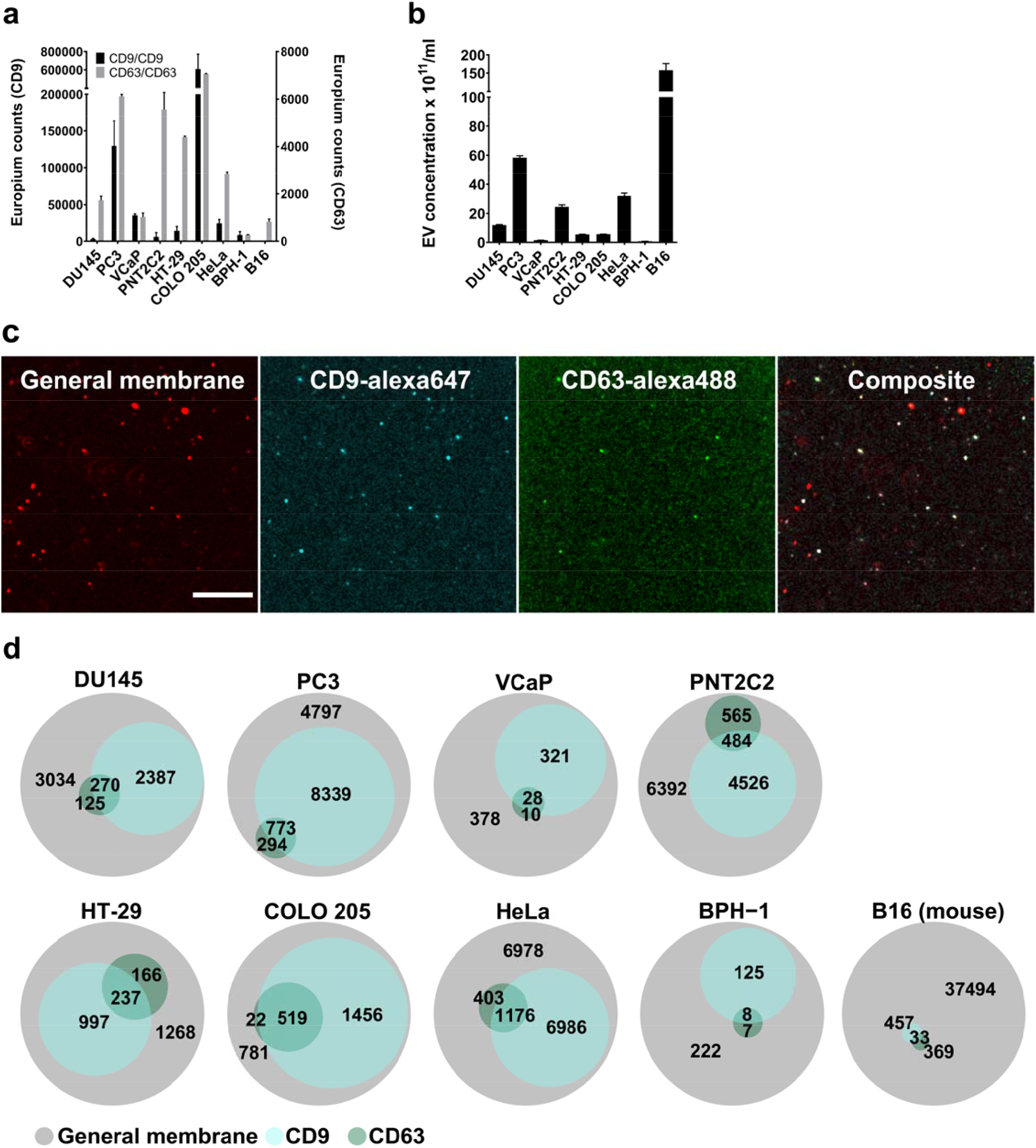
Quantification of EV subpopulations using immune specific detection. Isolated EVs from a panel of 9 cell lines (mouse B16 cell line as negative control) quantified by a human-specific CD9 and CD63 time-resolved immunoassay (TR-FIA) **(a)** and EVQuant **(b). (c)** Representative images (LSM510, Zeiss) of EVQuant analysis of fluorescently labeled EVs from a colon cancer cell line (COLO 205), using non-specific membrane labeling and specific immunofluorescent labeling of human CD9 and CD63. Scale bar 10 μm. Individual colour channels in the “Composite” image were slightly adjusted in terms of brightness/contrast to improve the visibility of the three different channels. **(d)** Venn Diagrams showing the different EV subpopulations detected in EV isolates from 9 different cell lines (B16 mouse cell line as negative control) using non-specific membrane labeling and specific immunofluorescent labeling of human CD9 and CD63. Images were acquired using a high-content screening system (Opera Phenix, Perkin Elmer). TR-FIA was performed in triplicate and EVQuant was performed in duplicate (Mean +- SD).

Together, these data demonstrate that EVQuant could not only quantify the total EV number but also enables quantification and analysis of individual EVs harboring specific biomarker combinations on their surface.

### EV quantification in clinical samples

There is a clear need in both research and clinical setting for efficient EV analysis in minimally processed biofluids. EV quantification in minimally processed urine showed a linear relation between measured concentrations and dilution (Supplemental Figure S5a). Analysis of 10 minimally processed urine samples showed a strong correlation between EVQuant and NTA, but again 1.9-fold higher EV concentrations by EVQuant compared to NTA (Figure 4a). As expected based on our previous work^12^, EVQuant analysis of urines collected from men after digital rectal examination (DRE) (n=10) showed significantly higher urinary EV concentrations compared to urines collected without DRE (n=10), after radical prostatectomy (n=10) and urines from women (n=10) (Figure 4b).

**Figure 4.**
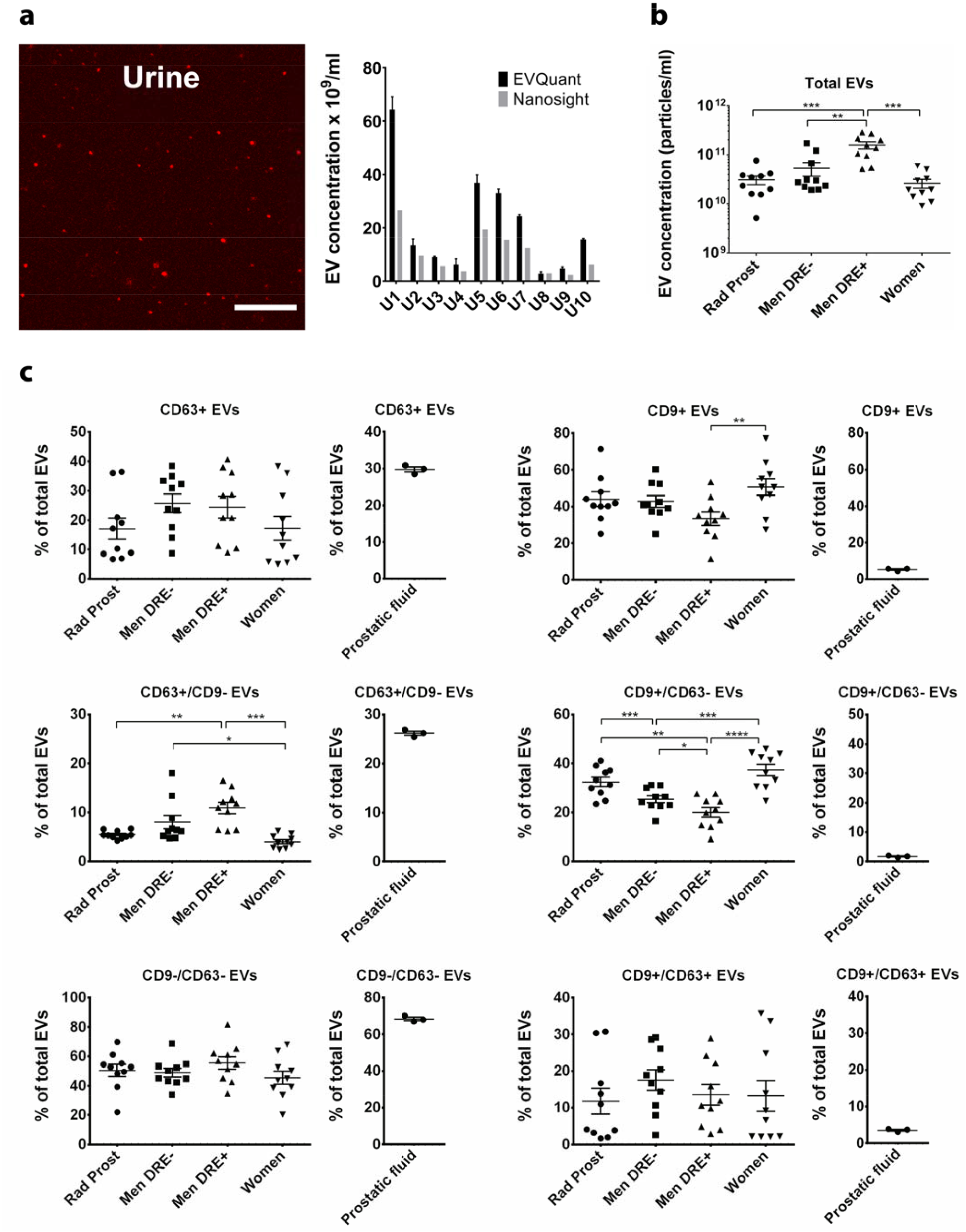
EV quantification in clinical urine samples. **(a)** EV quantification of ten minimally processed urine samples by EVQuant and NTA (right). Representative images of detected EVs in minimally processed urine determined by EVQuant (left). **(b)** EV concentration of a total of 40 minimally processed urines from men with and without a prior digital rectal examination (DRE), men that had a radical prostatectomy (Rad Prost) and women, quantified by EVQuant (Mean +- SEM). **(c)** EVQuant analysis of 40 minimally processed urines using a combination of non-specific membrane labeling and immune-specific labeling with human CD9 and CD63 immunofluorescent antibodies allowing relative quantification of specific EV subpopulations compared to total EVs (Mean +- SEM). Single NTA experiment was performed and EVQuant was performed in duplicate (Mean +- SD). For each comparison a Mann-Whitney rank sum test was performed. P-values: *0.05, **0.01, ***0.001, ****0.0001

As a proof-of-principle for EV biomarker analysis in clinical samples, we quantified sub-populations in urine supernatants based on CD9 and CD63 surface markers (Figure 4c, Supplemental Figure S5b). Interestingly, the fraction of CD9-negative and CD63-positive EVs (CD9-/CD63+) was significantly larger in healthy men, especially after receiving a DRE, and almost absent in women and men after radical prostatectomy, suggesting that most (CD9-/CD63+) EVs in urine are prostate-derived. We confirmed this in prostatic fluid from prostates obtained by radical prostatectomy^18^, where the CD9-/CD63+ fraction was the major fraction of the prostate-derived EVs (besides the double negative EVs) (Figure 4c). In contrast to most common techniques, EVQuant is also able to quantify EVs in minimally processed serum and plasma using immune detection of biomarkers present on the surface of EVs (e.g. CD9 and/or CD63) (Supplemental Figure S6a-b). The general EV labeling with Rhodamine, however, was not able to distinguish EVs from the high concentration of lipoproteins (Supplemental Figure S6a).

We show that EVQuant is able to quantify total EVs but also specific EV subpopulations in clinical samples such as minimally processed urine opening up new possibilities in both research and clinical settings.

### EV size determination by EVQuant

Fluorescent membrane labeling intensity increases quadratic with EV size and this relation could be applied to determine EV size. However, EV intensity a single plane is strongly affected by the positioning of EVs relative to the focal plane, resulting in dimmer, out of focus EVs which interfere with direct sizing based on fluorescence intensity in a single plane. This focus effect can be overcome by acquiring Z-stacks using small intervals and determining the maximum fluorescent intensity of the center of the point spread function (PSF) of each detected particle in 3D. As proof-of-concept, we analyzed monodispersed red fluorescent beads of different sizes (0.1, 0.2 and 0.5 μm) (Figure 5a). Unlike the typical EV labeling, fluorescence in beads is present in their volume, and thus intensity increases in diameter by the power third. The 100 nm beads’ fluorescent intensities were fitted using a gaussian function and mean intensity was used to calculate a calibration factor for converting fluorescent intensity to EV size (Figure 5b). In contrast to beads, the typical EV labeling results in membrane surface fluorescence rather than fluorescence in the lumen making liposomes more suitable for size calibration. We performed size analysis on liposomes having approximate sizes of 109, 44, and 35 nm (LIPO3, LIPO4 and LIPO5) as determined by DLS (Supplemental Table S4). EVQuant analysis was able to detect all three liposome types and showed that the intensity was related to liposome size (Figure 5c-d). The histograms show size distributions in line with the liposome sizes determined by DLS (Figure 5d). Analysis of conditioned medium and EV isolates from the DU145 PCa cell line showed that these samples contained a large number of small EVs with a size around 30-40 nm but also larger EVs up to 200 nm in size (Figure 5e-f).

**Figure 5.**
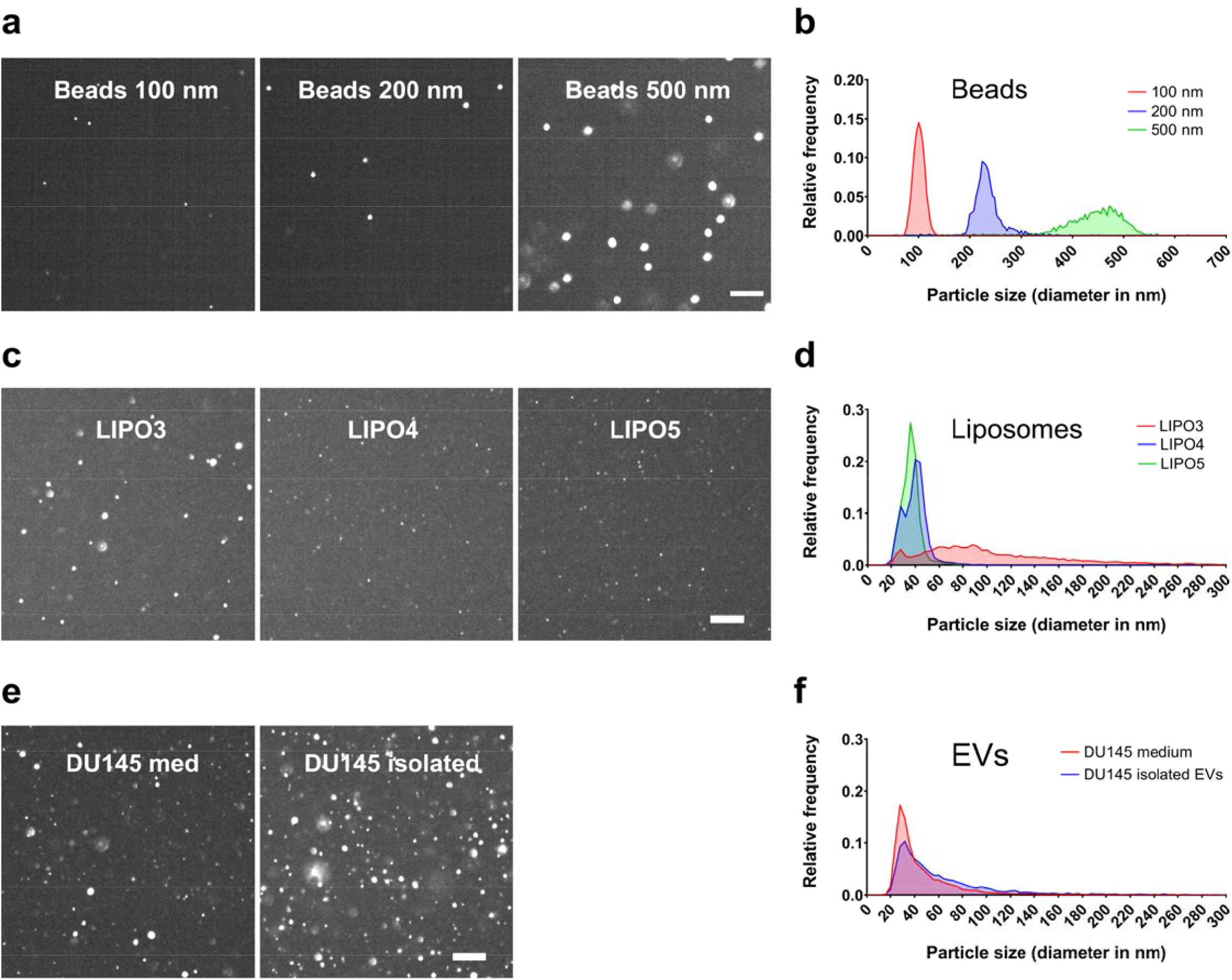
EV size determination by EVQuant by acquiring zstacks and performing 3D particle analysis. **(a)** Representative images of detected red fluorescent beads (FluoSpheres™, Invitrogen) of three different sizes (100, 200, and 500 nm) acquired by EVQuant. Scale bar 10 μm. **(b)** Size distributions of 100, 200 and 500 nm beads determined by EVQuant (100 nm beads were used as calibration particles). **(c)** Representative images of detected liposomes of three different sizes (109, 44 and 35 nm as determined by DLS) acquired by EVQuant. Scale bar 10 μm **(d)** Size distributions of liposomes of three different sizes determined by EVQuant (35 nm liposomes were used as calibration particles). **(e)** Representative images of detected EVs in minimally processed conditioned medium and EV isolate of prostate cancer cells (DU145) determined by EVQuant. **(f)** Size distributions of EVs in minimally processed conditioned medium and EV isolate of DU145 cells determined by EVQuant (35 nm liposomes were used as calibration particles).

Together, this data shows that EVQuant besides EV quantification also allows EV size characterization of small EVs at least down to 35 nm in size, corroborating the detection of the smallest liposomes in Figure 2.

## Discussion

EVs are a promising source of biomarkers and their abundance in body fluids makes them ideal for minimally invasive diagnostic, prognostic and monitoring assays. This is emphasized by the large number of studies focusing on discovery of novel EV-related biomarkers for a wide range of diseases and the many initiatives to develop technologies and assays to quantify EVs and EV-related protein/RNA biomarkers^4, 15, 19-23^. However, EV analysis in complex biological and clinical samples has shown to be a major challenge^24, 25^ and shows the need for novel assays using a simple, fast, sensitive and accurate technology in combination with standardized procedures for sample collection, sample treatment and analysis^15, 26^.

To address these challenges, we developed a straightforward and rapid EV quantification and characterization assay that allows multi-parameter analysis of individual EVs in biological fluids (EVQuant). The unique capture independent approach to immobilize EVs gives the EVQuant assay a major advantage, by allowing longer exposure times and therefore detection of low fluorescent signals. In addition, EV analysis is performed without extensive sample processing (isolation/purification) or EV capture procedures, thereby avoiding loss of EVs, underestimation of EV concentrations and biased analysis of EV subsets. Second, the EVQuant protocol (from sample preparation to EV-subtype analysis) only takes up a maximum of three hours while many other single EV analysis techniques report laborious protocols taking many hours^27–30^. Third, the EVQuant assay can be performed on conventional, widely available laser scanning confocal microscopes as well as on high-content screening systems for high-throughput analysis of up to 60 samples per hour. EVs are reliably detected over a broad range of concentrations (1E9-1E11 particles/ml) with a limit of detection of ~5E8 particles/ml, covering most clinical samples without the need for dilution or concentration of samples (Figure 1, Supplemental Figure S3a, S5a). Fourth, EVQuant showed ~2 to 4-fold higher particle concentration in general and even a ~5.5-fold higher concentration of small (~40 nm) liposomes than NTA analysis, suggesting a more sensitive detection of (small) particles in the EVQuant assay. This improved sensitivity is explained by the immobilization of EVs, allowing longer exposure times and detection of low intensity signals. This difference mostly influences the detection of smaller vesicles, as they intrinsically have lower signals and move faster leading to a lower chance to meet the inclusion requirements for NTA analysis.

Moreover, EVQuant is also able to provide particle size distributions using a 3D EV intensity analysis. Current EV quantification and characterization approaches are often limited by efficient detection of primarily larger vesicles. For example, NTA has an estimated limit for efficient detection of EVs of 60-70 nm^31^ while dedicated high-resolution flow cytometry and imaging flow cytometry are able to efficiently detect EVs as small as 60-100 nm^27, 28, 30, 32-35^. EVQuant size analysis is based on 3D intensity analysis of individual particles in the z-stack. Analysis showed a clear difference between size-distributions of liposomes of 35 nm and 109 nm in size (Figure 5D) and closely matched the size distributions as determined by dynamic light scattering (DLS) (Figure 5e). Equally important, this shows that EVQuant has an EV size detection limit of at least 35 nm, which is much lower compared to most current EV quantification methods. This shows to be highly relevant, as the majority of EVs in both the conditioned medium and isolated EVs from DU145 prostate cancer cells are smaller than 60 nm and are therefore missed by the majority of EV analysis approaches^31, 36^. It should however be noted that accurate size analysis by EVQuant is depended on the quality of the calibration sample. The accuracy could be hampered by heterogeneity in size of calibration particles, emphasizing the need for monodispersed membrane particles (e.g. liposomes) for calibration of current and newly developed EV analysis technologies.

Another important advantage of EVQuant is the ability to identify EV subpopulations by detection of multiple immunofluorescent markers on the surface of individual EVs. Interestingly, detection of the ‘general’ EV markers CD9 and CD63 on cell line-derived EVs showed that 37 to 71% were positive for CD9 and only 4 to 20% were CD63 positive. The differences in the presence of CD9 and CD63 are most likely related to relative expression of the markers in these cell lines^37^ or can be due to differences in the CD63 and CD9 antibody affinities. More importantly, this indicates that while CD9 and CD63 are often used as general EV markers, they are only detected on a fraction of EVs. EVQuant also provides the antibody signal intensity of individual EVs which can be used as an indication for the relative number of epitopes on an EV and allows to study differences in levels of biomarkers between samples.

Characterization of EV subpopulations in clinical urine samples identified prostate-derived EVs to be mainly negative for CD9 and positive for CD63 (CD9-/ CD63+). This prostate-derived EV marker profile was confirmed in prostatic fluid. Cell and tissue specific presence of these ‘general’ EV markers will have important consequences for (immune capture-based) assays that are dependent on these surface markers. It must be noted that the number of marker molecules on the surface of individual EVs required for EVQuant detection is currently unknown and could limit the application to detect the probably less abundant disease related markers. Improvements in fluorescent labeling of EVs like quantum dot labeling or signal amplification strategies like used in the proximity ligation assay (PLA)^38^ will increase the sensitivity of the EVQuant assay even further.

For two other important liquid biopsy sources, serum and plasma, EVQuant is able to quantify EVs without any laborious EV isolation or purification procedures. As with many EV assays, total EV quantification based on non-specific labeling is hampered by the high lipoprotein levels in serum and plasma, but the antibody-based detection still allows quantification of EVs in serum and plasma. Future development of novel fluorescent dyes or the identification of a novel biomarker which is able to discriminate EVs from lipoproteins, might improve quantification of total EVs in complex clinical samples like plasma and serum.

In summary, EVQuant is a rapid, sensitive and high throughput assay that allows quantification and sizing of individual EVs in several synthetic, biological and complex clinical samples without extensive isolation/purification procedures, limiting the loss of EVs and processing time. The ability to detect individual EVs down to 35 nm in size and the analysis of multiple biomarkers opens up many opportunities for the identification and quantification of novel specific EV subpopulations previously missed by bulk measurements in both research and clinical samples. Newly identified disease markers on the EV surface can be implemented in the EVQuant assay, resulting in a versatile assay that is easily applicable for a range of diseases including urogenital cancers. Together, this demonstrates an important application for the EVQuant assay in a clinical setting, where it can be used to detect organ- or disease-specific EVs.

## Acknowledgements

This work was supported by the IMMPROVE Alpe d’HuZes grant of the Dutch Cancer Society (EMCR2015-8022), PROSCANEXO TRANSCAN (JTC2016) sponsored by the Dutch Cancer Society (EMCR2017-8304), Prostate Cancer UK grant (G2012-36), the CANCER-ID Netherlands Organisation for Scientific Research-Domain Applied and Engineering Sciences (NWO-TTW) research program 14194, and the Daniel den Hoed Foundation grant for Erasmus MC Cancer Treatment Screening Facility. RvdM is supported by a VENI Fellowship (# 14385) from the Netherlands Organization for Scientific Research (NWO).

## Author contributions

MEvR, GWJ and TAH conceived the ideas and designed the study, TAH, ND, MSV and MEvR performed the experiments and analyzed the data, DD and GWJ supported the TR-FIA analyses, RvdM, RMS supported the synthetic vesicles/ liposomes work, PJF, WvC and ABH developed and supported with the imaging platforms, TAH and JAS wrote the script for image analysis, JAK performed TEM, MEvR, ABH and GWJ supervised the study and interpreted the data, TAH and MEvR wrote the manuscript.

## Declaration of Interest Statement

Conflict of interest: GWJ has a non-exclusive license agreement with Cell Guidance Systems for the TR-FIA assay.

## Data availability statement

The data that support the findings of this study are available from the corresponding author upon reasonable request.

## Code availability statement

The EVQuant source code and all other corresponding data (e.g. plugin, demo data, installation guide, license) can be found in the following open source repository upon acceptance of the publication: https://github.com/MEvanRoyen/EVQuant. The EVQuant plugin is licensed under a GNU General Public License v3 (GPLv3).

## Notes

https://github.com/MEvanRoyen/EVQuant

## References

1. Thery, C. et al. Minimal information for studies of extracellular vesicles 2018 (MISEV2018): a position statement of the International Society for Extracellular Vesicles and update of the MISEV2014 guidelines. J Extracell Vesicles 7, 1535750 (2018).

2. Stoorvogel, W., Kleijmeer, M.J., Geuze, H.J. & Raposo, G. The biogenesis and functions of exosomes. Traffic 3, 321–330 (2002).

3. Duijvesz, D., Luider, T., Bangma, C.H. & Jenster, G. Exosomes as biomarker treasure chests for prostate cancer. European urology 59, 823–831 (2011).

4. Skog, J. et al. Glioblastoma microvesicles transport RNA and proteins that promote tumour growth and provide diagnostic biomarkers. Nature cell biology 10, 1470–1476 (2008).

5. Junker, K., Heinzelmann, J., Beckham, C., Ochiya, T. & Jenster, G. Extracellular Vesicles and Their Role in Urologic Malignancies. European urology (2016).

6. Merchant, M.L., Rood, I.M., Deegens, J.K.J. & Klein, J.B. Isolation and characterization of urinary extracellular vesicles: implications for biomarker discovery. Nat Rev Nephrol 13, 731–749 (2017).

7. Moon, P.G. et al. Identification of Developmental Endothelial Locus-1 on Circulating Extracellular Vesicles as a Novel Biomarker for Early Breast Cancer Detection. Clin Cancer Res 22, 1757–1766 (2016).

8. van der Pol, E. et al. Optical and non-optical methods for detection and characterization of microparticles and exosomes. Journal of thrombosis and haemostasis: JTH 8, 2596–2607 (2010).

9. Maas, S.L.N., De Vrij, J. & Broekman, M.L.D. Quantification and Size-profiling of Extracellular Vesicles Using Tunable Resistive Pulse Sensing. Jove-J Vis Exp (2014).

10. Erdbrugger, U. & Lannigan, J. Analytical challenges of extracellular vesicle detection: A comparison of different techniques. Cytometry. Part A: the journal of the International Society for Analytical Cytology 89, 123–134 (2016).

11. Jorgensen, M. et al. Extracellular Vesicle (EV) Array: microarray capturing of exosomes and other extracellular vesicles for multiplexed phenotyping. J Extracell Vesicles 2 (2013).

12. Duijvesz, D. et al. Immuno-based detection of extracellular vesicles in urine as diagnostic marker for prostate cancer. International journal of cancer 137, 2869–2878 (2015).

13. Zarovni, N. et al. Integrated isolation and quantitative analysis of exosome shuttled proteins and nucleic acids using immunocapture approaches. Methods 87, 46–58 (2015).

14. Kanwar, S.S., Dunlay, C.J., Simeone, D.M. & Nagrath, S. Microfluidic device (ExoChip) for on-chip isolation, quantification and characterization of circulating exosomes. Lab Chip 14, 1891–1900 (2014).

15. Hartjes, T.A., Mytnyk, S., Jenster, G.W., van Steijn, V. & van Royen, M.E. Extracellular Vesicle Quantification and Characterization: Common Methods and Emerging Approaches. Bioengineering (Basel) 6 (2019).

16. Maas, S.L. et al. Possibilities and limitations of current technologies for quantification of biological extracellular vesicles and synthetic mimics. J Control Release 200, 87–96 (2015).

17. van der Pol, E. et al. Particle size distribution of exosomes and microvesicles determined by transmission electron microscopy, flow cytometry, nanoparticle tracking analasis, and resistive pulse sensing. Journal of thrombosis and haemostasis: JTH 12, 1182–1192 (2014).

18. Fujita, K. et al. Cytokine profiling of prostatic fluid from cancerous prostate glands identifies cytokines associated with extent of tumor and inflammation. Prostate 68, 872–882 (2008).

19. Del Re, M. et al. The Detection of Androgen Receptor Splice Variant 7 in Plasma-derived Exosomal RNA Strongly Predicts Resistance to Hormonal Therapy in Metastatic Prostate Cancer Patients. European urology 71, 680–687 (2017).

20. Choi, D.S. Urinary extracellular vesicles for biomarker source to monitor polycystic kidney disease. Proteomics. Clinical applications 9, 447–448 (2015).

21. Etayash, H., McGee, A.R., Kaur, K. & Thundat, T. Nanomechanical sandwich assay for multiple cancer biomarkers in breast cancer cell-derived exosomes. Nanoscale 8, 15137–15141 (2016).

22. Jakobsen, K.R. et al. Exosomal proteins as potential diagnostic markers in advanced non-small cell lung carcinoma. Journal of Extracellular Vesicles 4 (2015).

23. Minciacchi, V.R., Zijlstra, A., Rubin, M.A. & Di Vizio, D. Extracellular vesicles for liquid biopsy in prostate cancer: where are we and where are we headed? Prostate cancer and prostatic diseases 20, 251–258 (2017).

24. Ramirez, M.I. et al. Technical challenges of working with extracellular vesicles. Nanoscale 10, 881–906 (2018).

25. Lane, R.E., Korbie, D., Hill, M.M. & Trau, M. Extracellular vesicles as circulating cancer biomarkers: opportunities and challenges. Clin Transl Med 7 (2018).

26. Witwer, K.W. et al. Standardization of sample collection, isolation and analysis methods in extracellular vesicle research. J Extracell Vesicles 2 (2013).

27. Gorgens, A. et al. Optimisation of imaging flow cytometry for the analysis of single extracellular vesicles by using fluorescence-tagged vesicles as biological reference material. J Extracell Vesicles 8, 1587567 (2019).

28. van der Vlist, E.J., Nolte-’t Hoen, E.N.M., Stoorvogel, W., Arkesteijn, G.J.A. & Wauben, M.H.M. Fluorescent labeling of nano-sized vesicles released by cells and subsequent quantitative and qualitative analysis by high-resolution flow cytometry. Nat Protoc 7, 1311–1326 (2012).

29. Lee, K. et al. Multiplexed Profiling of Single Extracellular Vesicles. ACS nano 12, 494–503 (2018).

30. Ricklefs, F.L. et al. Imaging flow cytometry facilitates multiparametric characterization of extracellular vesicles in malignant brain tumours. J Extracell Vesicles 8, 1588555 (2019).

31. Bachurski, D. et al. Extracellular vesicle measurements with nanoparticle tracking analysis - An accuracy and repeatability comparison between NanoSight NS300 and ZetaView. Journal of Extracellular Vesicles 8 (2019).

32. Stoner, S.A. et al. High sensitivity flow cytometry of membrane vesicles. Cytometry. Part A: the journal of the International Society for Analytical Cytology 89, 196–206 (2016).

33. Pospichalova, V. et al. Simplified protocol for flow cytometry analysis of fluorescently labeled exosomes and microvesicles using dedicated flow cytometer. Journal of Extracellular Vesicles 4 (2015).

34. de Rond, L. et al. A Systematic Approach to Improve Scatter Sensitivity of a Flow Cytometer for Detection of Extracellular Vesicles. Cytometry. Part A: the journal of the International Society for Analytical Cytology (2020).

35. Zhu, S. et al. Light-scattering detection below the level of single fluorescent molecules for high-resolution characterization of functional nanoparticles. ACS nano 8, 10998–11006 (2014).

36. Puhka, M. et al. Metabolomic Profiling of Extracellular Vesicles and Alternative Normalization Methods Reveal Enriched Metabolites and Strategies to Study Prostate Cancer-Related Changes. Theranostics 7, 3824–3841 (2017).

37. Koliha, N. et al. A novel multiplex bead-based platform highlights the diversity of extracellular vesicles. Journal of Extracellular Vesicles 5 (2016).

38. Lof, L. et al. Detection of Extracellular Vesicles Using Proximity Ligation Assay with Flow Cytometry Readout-ExoPLA. Current protocols in cytometry 81, 4 8 1–4 8 10 (2017).

39. Duijvesz, D. et al. Proteomic profiling of exosomes leads to the identification of novel biomarkers for prostate cancer. PloS one 8, e82589 (2013).

40. Kulkarni, J.A. et al. On the Formation and Morphology of Lipid Nanoparticles Containing Ionizable Cationic Lipids and siRNA. ACS nano 12, 4787–4795 (2018).

41. Schindelin, J. et al. Fiji: an open-source platform for biological-image analysis. Nature methods 9, 676–682 (2012).

42. Duijvesz, D. et al. Immuno-based detection of extracellular vesicles in urine as diagnostic marker for prostate cancer. International journal of cancer 137, 2869–2878 (2015).

43. Kulkarni, J.A. et al. Fusion-dependent formation of lipid nanoparticles containing macromolecular payloads. Nanoscale 11, 9023–9031 (2019).

