## Supplemental Data and Information for "EVQuant; high-throughput quantification and characterization of extracellular vesicle (sub)populations"

#### Legends supplementary figures

**Supplemental Figure S1. Calibration of the imaging volume of a laser scanning confocal microscope (LSM510, Zeiss) to allow absolute particle quantification (a)** 3D visualization of a 10 µm z-stack (interval 0.5 µm) shows the point-spread function of the fluorescent particles in z-direction which results in detection of the same particle in multiple slices (left)**.** XY and corresponding YZ orthogonal view of the z-stack (right). Scale bar 10 µm. **(B)** Serial dilutions of 100 nm tetraspeck beads are quantified using EVQuant and measured concentrations are plotted against calculated concentrations based on specifications given by the manufacturer (right). Representative image of detected 100 nm tetraspeck beads acquired by EVQuant (left). Scale bar 5 µm. EVQuant was performed in duplicate (Mean +- SD).

**Supplemental Figure S2. Characterization of membrane nanoparticles. (a)** Particle size of LIPO1 and LIPO2 lipid nanoparticles obtained by NTA (mode), DLS (number) and EM (median). **(b)** Size distributions of LIPO1 and LIPO2 lipid nanoparticles based on analysis of multiple images acquired by Cryo-EM (~100 particles per sample, bin size 4 nm).

**Supplemental Figure S3. EV quantification in biological samples. (a)** Serial dilutions of minimally processed cell conditioned medium from prostate cancer cells (VCaP) quantified by EVQuant. **(b-c)** EV concentration of minimally processed cell conditioned medium and matching EV isolates of a panel of 10 cell lines from different origins was quantified by EVQuant and NTA (NanoSight NS300). Measured concentrations by EVQuant are plotted against measured concentrations by NTA. **(d)** Calculated EV recovery after EV isolation using a standard ultra-centrifugation protocol. **(e)** Comparison of EVQuant analysis using two different microscopy-based systems. EV concentrations of isolated EVs measured by a laser scanning confocal microscope (LSM510, Zeiss) were plotted against concentrations measured by a High Content Screening System (Opera Phenix™, Perkin Elmer). Single experiment was performed for NTA and EVQuant assay was performed in duplicate (Mean +- SD).

**Supplemental Figure S4. EV quantification of isolated EVs from a panel of 9 cell lines (mouse B16 cell line as negative control) using a human-specific CD9 and CD63 time-resolved immunoassay (TR-FIA) and EVQuant by combining non-specific membrane labeling and immune-specific labeling using human CD9 and CD63 immunofluorescent antibodies. (a)** Measured europium counts by a CD9 and CD63 TR-FIA are plotted against each other. Measured CD9 **(b)** and CD63 **(c)** europium counts plotted against total EV concentrations measured by EVQuant. **(d)** The effect of immune-specific antibodies on EVQuant concentration measurements was determined by quantification of isolated EVs from a prostate cancer cell line (DU145) with and without the presence of CD9 or CD63 immune-specific antibodies. **(e-f)** CD9 and CD63 europium counts measured by TR-FIA divided by concentration of CD9+ and CD63+ EVs measured by EVQuant. **(g)** The median CD9 and CD63 intensity of detected CD9+ and CD63+ EVs is determined for each cell line as a measure for the relative number of biomarker epitopes present on the EVs. **(h-i)** CD9 and CD63 europium counts (normalized for the number of CD9+ and CD63+ EVs) were plotted against the median CD9 and CD63 spot intensities.

**Supplemental Figure S5. Quantification of EV subpopulations in clinical urine samples. (a)** Serial dilutions of minimally processed urine from a healthy volunteer were quantified by EVQuant. **(b)** Absolute quantification of specific EV subpopulations in 40 urines by combining non-specific and immune-specific labeling using CD9-alexa647 and CD63-alexa488 immunofluorescent antibodies (Mean +- SEM). EVQuant assay was performed in duplicate (Mean +- SD).

**Supplemental Figure S6. EV quantification in blood (serum and plasma). (a)** Representative images of EV subpopulations in serum from a prostate cancer patient acquired by EVQuant by combining non-specific membrane labeling and immune-specific labeling using CD9 and CD63 immunofluorescent antibodies. **(b)** Quantification of EV subpopulations in serum and plasma samples from 5 prostate cancer patients based on immune-specific labeling using CD9 and CD63 immunofluorescent antibodies.

#### Supplemental Figure S1

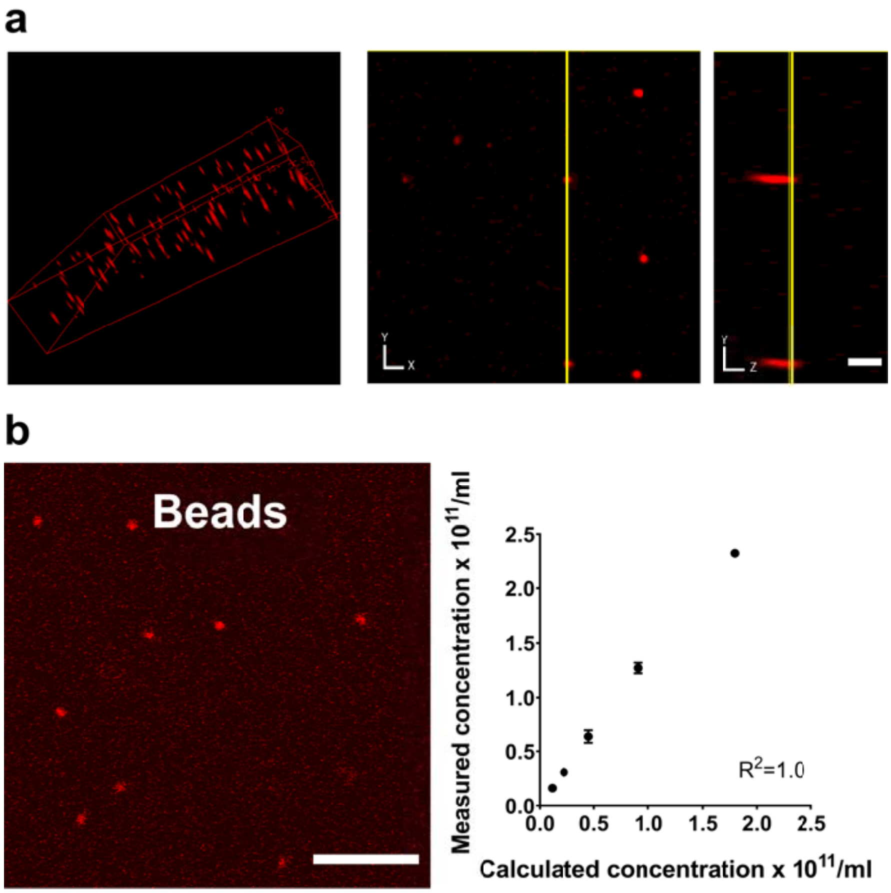

### Supplemental Figure S2

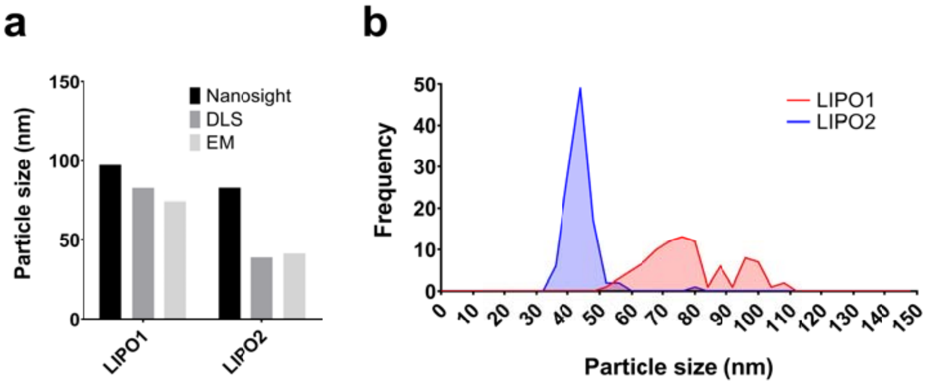

### Supplemental Figure S3

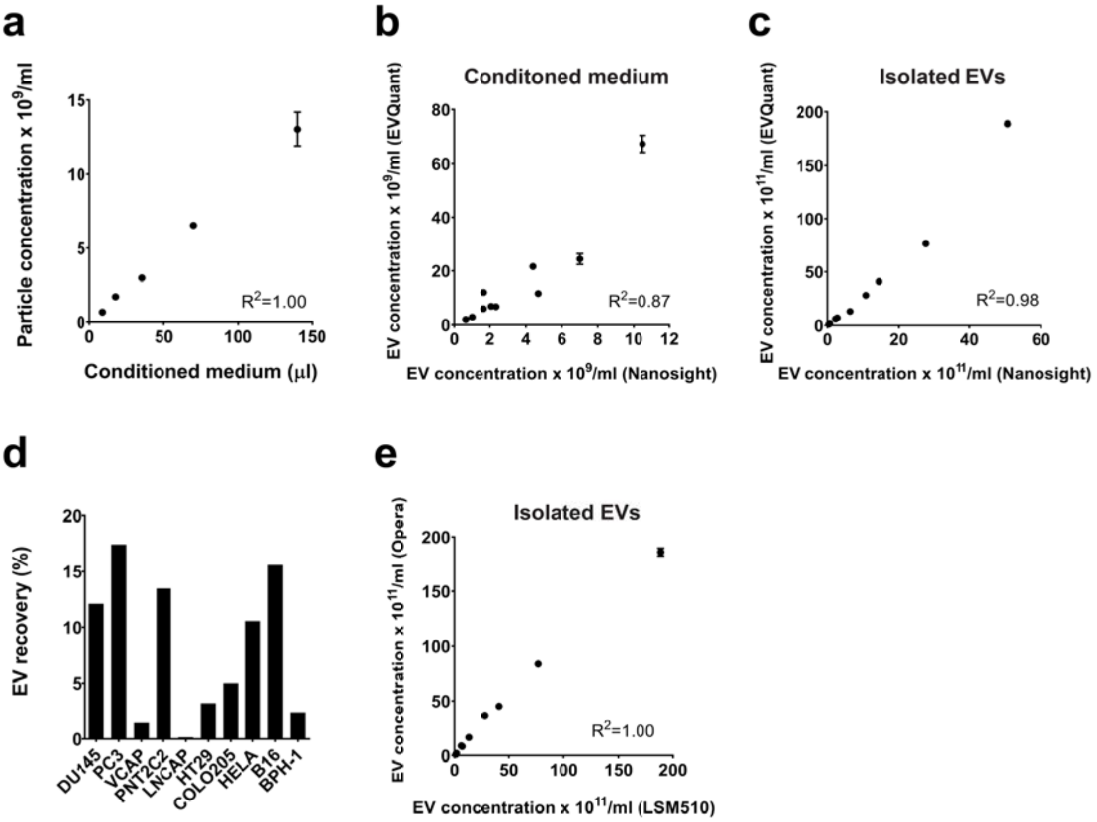

### Supplemental Figure S4

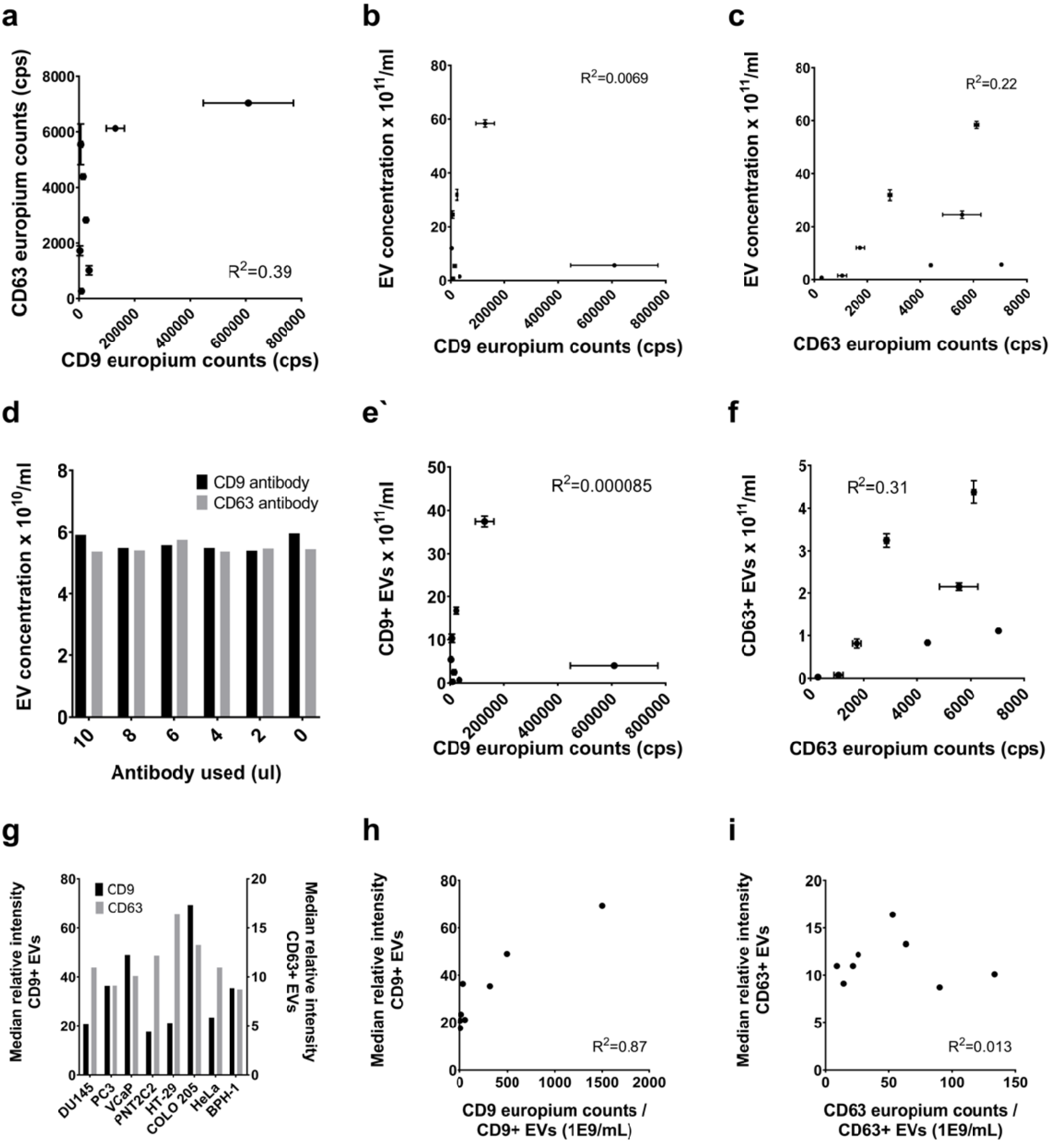

### Supplemental Figure S5

**a**

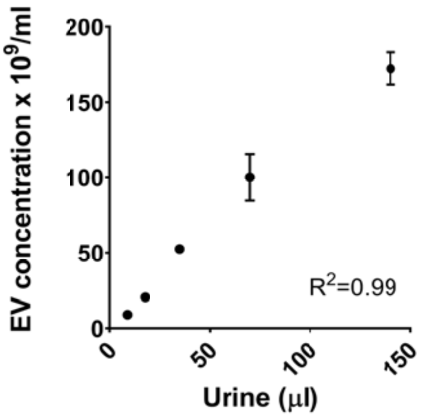


**b**

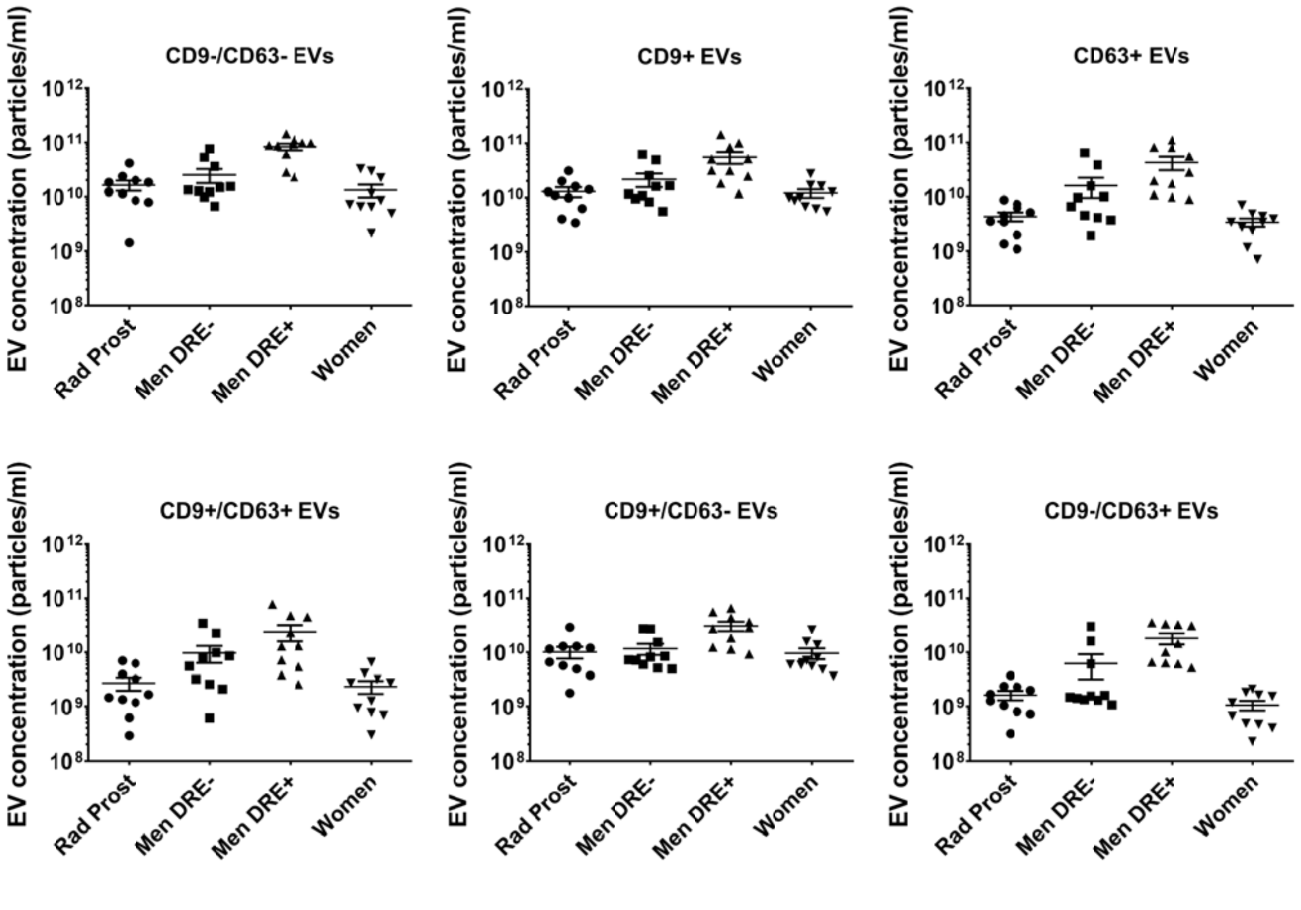

### Supplemental Figure S6

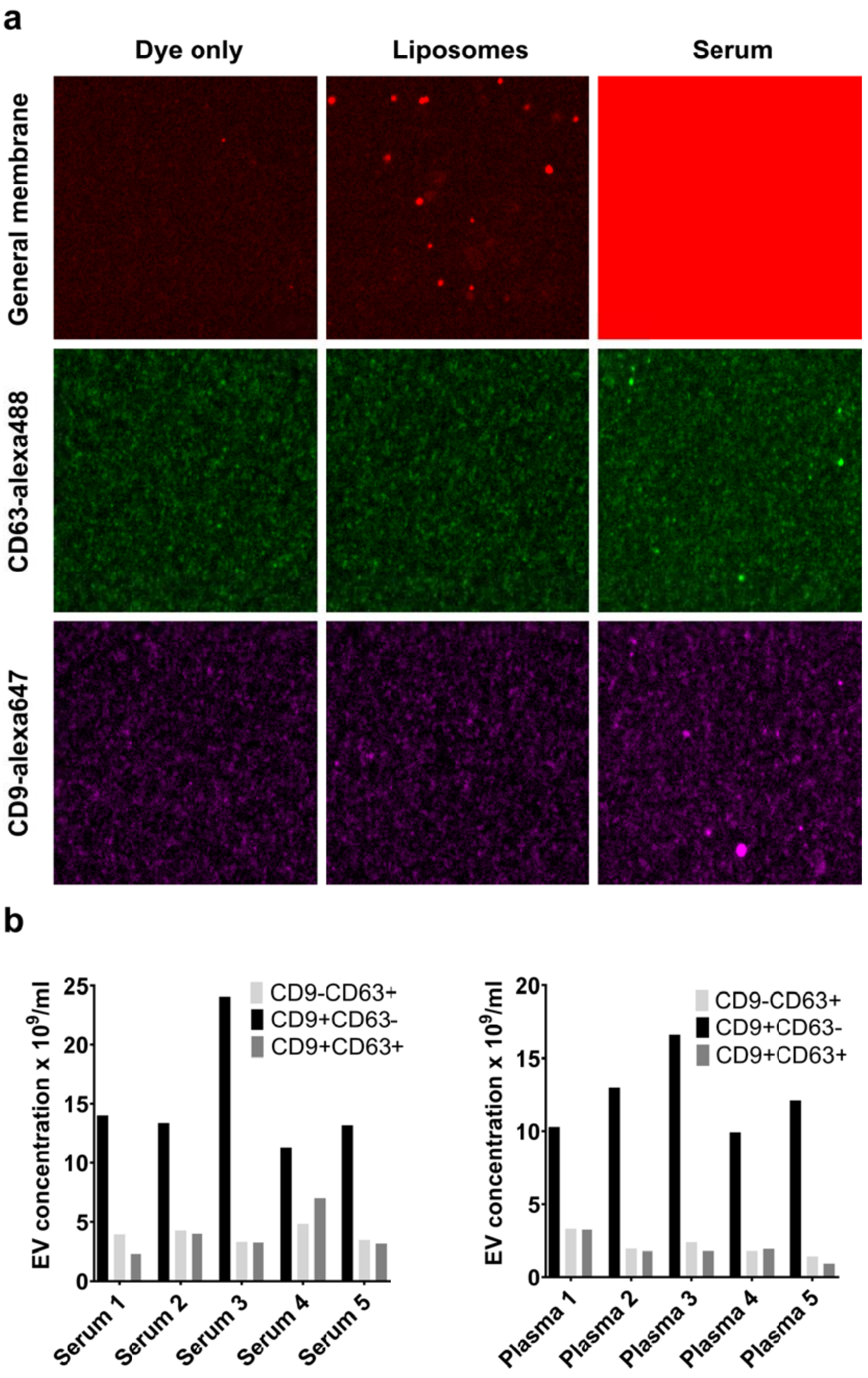

**Supplemental Table S1: Effective optical slice tickness LSM510 (Zeiss)**

| **Sample:** | **DU145 medium** |
| --- | --- |
| Microscope: | LSM510 Laser scanning confocal micrsocope (Zeiss) |
| Detection threshold imageJ: | 4 |
| Laser output (543nm): | 4.53 uW |
| Number of stacks measured: | 4 |
| Average EOST: | 1.673 um SD: 0.107 |

| **Sample:** | **Tetraspeck beads 0.1 um diameter** |
| --- | --- |
| Microscope: | LSM510 Laser scanning confocal micrsocope (Zeiss) |
| Detection threshold imageJ: | 4 |
| Laser output (543nm): | 4.53 uW |
| Number of stacks measured: | 4 |
| Average EOST: | 1.688 SD: 0.1 |

**Supplemental Table S2. Characterization of LNPs by DLS and Cholesterol E assay (Batch 1)**

| | | | Cholesterol E assay | | DLS (mean) | | | | | | | | | |
|---|---|---|---|---|---|---|---|---|---|---|---|---|---|---|
| # | DSPC/Cholesterol/DSPE-PEG2000 | Microfluidics flow ratio (O:A) | [lipid mg/mL] | [lipid mM] | Z-ave | SD | PDI | SD | Intensity | SD | Number | SD | Volume | SD |
| LNP1 | 53/42/5 | 1:1 | 5.47 | 7.58 | 128.37 | 1.80 | 0.16 | 0.02 | 181.02 | 45.29 | 82.61 | 6.42 | -19.93 | 0.64 |
| LNP2 | 53/42/5 | 1:3 | 6.31 | 8.74 | 54.69 | 0.69 | 0.10 | 0.03 | 59.56 | 2.85 | 39.00 | 1.71 | -18.63 | 1.72 |

**Supplemental Table S3. Effective optical slice tickness Opera Phenix High Content Screening Microscope (Perkin Elmer)**

| **Sample:** | **DU145 medium** |
|---|---|
| Microscope: | Opera Phenix High Content Screening Microscope (Perkin Elmer) |
| Detection threshold imageJ: | 19 |
| Laser output (568nm): | 100% |
| Exposure time: | 500 msec |
| Number of stacks measured: | 4 |
| Average EOST: | 2.172 um SD: 0.018 |

| **Sample:** | **Tetraspeck beads 0.1 um diameter** |
|---|---|
| Microscope: | Opera Phenix High Content Screening Microscope (Perkin Elmer) |
| Detection threshold imageJ: | 19 |
| Laser output (568nm): | 100% |
| Exposure time: | 500 msec |
| Number of stacks measured: | 4 |
| Average EOST: | 3.232 SD: 0.01 |

**Supplemental Table S4. Characterization of LNPs by DLS and Cholesterol E assay Batch 2**

| | | | Phospholipid C assay | | DLS (mean) | | | | | | | | | |
|---|---|---|---|---|---|---|---|---|---|---|---|---|---|---|
| # | DSPC/Cholesterol/DSPE-PEG2000 | Microfluidics flow ratio (O:A) | [lipid mg/mL] | [lipid mM] | Z-ave | SD | PDI | SD | Intensity | SD | Number | SD | Volume | SD |
| LNP3 | 53/42/5 | 1:1 | 1.82 | 2.52 | 168.80 | 2.16 | 0.14 | 0.01 | 197.30 | 5.85 | 109.40 | 8.75 | 194.40 | 6.20 |
| LNP4 | 53/42/5 | 1:2 | 4.76 | 6.59 | 59.20 | 0.54 | 0.06 | 0.01 | 63.80 | 0.86 | 43.90 | 0.25 | 52.40 | 0.25 |
| LNP5 | 53/42/5 | 1:3 | 5.48 | 7.59 | 52.70 | 0.49 | 0.12 | 0.02 | 58.70 | 0.59 | 35.30 | 2.04 | 44.00 | 1.58 |

**Supplemental Table S5. Cell lines and culture media used to grow the cell[illegible]**

| Origin | Name | Species | Medium | FCS |
|---|---|---|---|---|
| Prostate cancer | DU145 | Human | RPMI 1640 | 5% |
| | PC3 | Human | RPMI 1640 | 5% |
| | VCaP | Human | RPMI 1640 | 10% |
| | LNCaP | Human | RPMI 1640 | 5% |
| Normal prostate | PNT2C2 | Human | RPMI 1640 | 5% |
| Benign prostate hyperplasia | BPH-1 | Human | RPMI 1640 | 10% |
| Colon cancer | COLO 205 | Human | RPMI 1640 | 10% |
| | HT-29 | Human | DMEM | 10% |
| Cervical cancer | HeLa | Human | DMEM | 10% |
| Skin cancer | B16 | Mouse | DMEM | 10% |

### EVQuant; high-throughput quantification and characterization of extracellular vesicle (sub)populations

### Table of contents

SUMMARY ... 3
MATERIALS ... 4
REAGENTS ... 4
EQUIPMENT ... 4
REAGENTS SETUP ... 6
PROCEDURES ... 7
1. EVQuant: Sample preparation ... 7
2. EVQuant: Setting up the microscope ... 9
3. EVQuant: Total EV quantification using non-specific membrane labeling ... 13
4. EVQuant: Quantification of EV subpopulations by combining general membrane labeling and immune-specific labeling ... 15
5. EVQuant: EV size characterization ... 19
APPENDIX ... 21

### SUMMARY

The EVQuant assay is developed to detect individual EVs to quantify the total number and/or size and/or biomarker specific subpopulations of extracellular vesicles in several biological fluids such as EV isolations, cell-conditioned media, plasma and serum. This assay is based on detection of fluorescent nanoparticles using a confocal microscope. EVs are either non-specifically labeled using the fluorescent membrane dye Octadecyl Rhodamine B Chloride R18 (Invitrogen, final staining concentration of 0.33 ng/ul) or PKH26/67 dye (Sigma-Aldrich, final staining concentration of 5μM) and/or specifically labeled using the immunofluorescent Mouse CD9 Monoclonal (MEM-61)–Alexa Fluor® 647 antibody (Invitrogen, 1:25 dilution, 50 μl staining volume) and/or Mouse CD63 monoclonal (MX-49.129.5)-Alexa Fluor®488 antibody (Santa Cruz, catalog# sc-5275 AF488, antibody concentration 2.5 μg/ml, 50 μl staining volume). Subsequently, the fluorescent nanoparticles are immobilized in a non-denaturing polyacrylamide gel (final ratio of 16% (w/w) acrylamide/bis-acrylamide) to enable longer exposure times and detection of low intensity signals. Images are acquired by confocal microscopy to minimize background fluorescence of free dye, using a conventional laser scanning confocal microscopy (e.g. LSM510Meta, Zeiss) or a high-content screening system (e.g. Opera Phenix, Perkin Elmer). Phosphate buffered saline (PBS) and/or serum free culture medium, supplemented with the same concentration of fluorescent dyes/antibodies are used as negative controls. Cell conditioned medium (DU145 prostate cancer cells), liposomes and/or 100 nm fluorescent tetraspeck beads (Invitrogen) are used as internal standards for quality control and system calibration. Images are analyzed by open source software ImageJ using the EVQuant plugin (latest version will be available upon acceptance for publication at the open source repository https://github.com/MEvanRoyen/EVQuant and further information can be found at http://www.evquant.nl) or the Opera Phenix analysis software (Harmony 5.4, Perkin Elmer). Several detailed stepwise protocols to prepare, calibrate, perform and analyze EVQuant measurements are provided below. A flow chart of the analysis procedures is presented in the Appendix (Figure 1).

### MATERIALS

#### REAGENTS

*Tissue culture*

- DU145 cells
- RPMI 1640 with L-Glutamine (Lonza, cat. no. #Be12-702F)
- FBS (Lonza, cat. no. DE14-801F)
- Penicillin-streptomycin (pen/strep; 5,000 U/ml (Lonza, cat. no. 17-603E)
- Phosphate Bufferd Saline (Lonza, cat. no. #17-516F)
- Trypsin-Versene (EDTA) Mixture (Lonza, cat. no. #BE17-161E)
- DSPC-PEG-DSPE liposomes (100nm)

*Fluorescence*

- Mouse CD9 Monoclonal (MEM-61) – Alexa Fluor® 647 antibody (Thermo Fisher, cat. no. #MA5-18154, 100 tests/400ul)
- Mouse CD63 (MX-49.129.5) - Alexa Fluor® 488 antibody (Santa Cruz, cat. no. sc-5275 AF488, 200µg/ml)
- Octadecyl Rhodamine B Chloride (R18) (Life Technologies, cat. no. #O-246)
- TetraSpeck™ Microspheres, 0.1 µm, fluorescent blue/green/orange/dark red (Life Technologies, cat. no. T7279)

*Chemicals*

- Acrylamide/Bis Solution 37.5:1 30% w/v (SERVA, cat. no. #10688.01)
- Ultra-pure Temed (Life Technologies, cat. no. #15524-010)
- Ammonium persulfate (Life Technologies, cat. no. #7727-54-0)
- Methanol
- Demineralized water
- Bovine Serum Albumin (Sigma-Aldrich)

#### EQUIPMENT

- Humidified incubator, 5% CO2
- Centrifuge
- Biosafety cabinet level II
- T175 culture flasks (Greiner Bio-One, cat. no. 661-160)
- Conical 50-ml tubes (Greiner Bio-One, cat. no. 227-661)
- Falcon 5-ml round-bottom polystyrene tubes (BD Falcon, cat. no. 352058)
- Syringe filter unit, 0.22 µm
- Syringe filter unit, 0.45 µm
- Syringe, 50 ml
- Tube, Thinwall, Ultra-Clear™, 38.5 mL, 25 x 89 mm (Beckman Coulter, cat. no. 344058)
- Tube, Thinwall, Polypropylene, 4 mL, 11 x 60 mm (Beckman Coulter, cat. no. 328874)
- Ultracentrifuge + Rotors (swinging buckets) which allows centrifugation at 100,000xg (i.e. Beckman L-80 Optima Ultracentrifuge and SW32Ti + SW60 rotors)
- Greiner Sensoplate™ glass bottom 96-wells plate (Sigma-Aldrich, cat. no. M4187-16EA)
- CellVis P96-0-N glass bottom 96-wells imaging plate (CellVis, cat. No. P96-0-N)

- Confocal microscope having the right mirrors and filter sets for imaging with excitation wavelength around 540-560 nm and emission wavelength around 560-700 nm. Multicolour analysis, requires the microscope is suitable for additional excitation wavelengths around 490 and 640 nm and emission wavelengths of 500-550 and 650-700 nm, respectively (i.e. Zeiss, LSM 510 Axiovert 200M inverted microscope).
- 40x, NA 1.3, oil Plan-neofluar objective (or comparable objective)
- 488nm argon laser, HeNe 543 nm laser, HeNe 633 nm laser (other lasers that have a slightly different wavelength are also suitable)
- Build in macro software on the microscope for the acquisition of Tile-scans (optional)
- Computer that has the open source software Fiji and the EVQuant plugin installed (latest version will be available at the open source repository https://github.com/MEvanRoyen/EVQuant and further information can be found at http://www.evquant.nl).
- Perkin Elmer Opera Phenix equipped with appropriate lasers and filters
- Perkin Elmer Harmony operating and analysis software (version 4.9, but older versions will suffice)

### REAGENTS SETUP

**Complete culture medium**
Supplement RPMI 1640 L-Glutamine with 5% (vol/vol) FCS and add pen-strep to a final concentration of 50 U/ml penicillin and 50 μg/ml streptomycin. FCS and pen-strep are passed through 0.22 μm filter before addition to the medium. Store the medium for up to 3 months at 4°C.

**Bovine serum albumin (BSA) stock solution**
Dissolve 300 milligram of BSA powder in 100 ml PBS to get a 0.3% (wt/wt) BSA stock solution. Gently shaking the bottle will completely dissolve the BSA within several minutes. Filter the solution trough a 0.22 μm filter and make aliquots of 0.5 ml for long term storage at −80°C. Once BSA is thawed, it should be kept at 4°C and can be used for 1 week.

**Ammonium persulfate (APS), 10% (wt/vol)**
Dissolve 1 gram of APS powder in 10 ml demineralized water to get a 10% (wt/vol) APS stock solution. Make aliquots of 150 ul for long term storage at −80°C. Once APS is thawed, it should be kept at 4°C and used on the same day.

**Rhodamine working solution (25 ng/ul)**
Dissolve 25 μl of the Rhodamine stock solution (10 mg/ml) in 10 ml methanol to get a working solution of 25 ng/μl. Make aliquots of 1 ml, seal with paraffin membrane, and store the working solutions at −20°C. Working solutions can be stored at -20°C up to 6 months.

**Internal standards for EV quantification**
Plate out DU145 (or any other cell line) cells in a T175 culture flask and grow the cells in 25 ml complete culture medium. When cells reach 70-80% confluency, wash the cells twice with PBS and grow the cells in 25 ml FCS-free culture medium for 48 hours. Collect the conditioned medium in a 50 ml tube, centrifuge at 3000xg for 20 min at 4°C to remove cells and cell debris and collect the supernatant in a new 50 ml tube. Make aliquots of 40 ul for long term storage at −80°C (~500 standard samples are made using 20 ml of conditioned medium)

### PROCEDURES

#### 1. EVQuant: Sample preparation

1. 30 minutes before sample preparation, take out the 0.3% BSA and 10% APS from the -80 and -20°C, respectively, and thaw the vials at RT. Take a vial of Rhodamine working solution (25 ng/ul) from the -80°C and keep on ice.
2. Open the Excel template; "EVQuant_Template.xlsx" if EVs are only labeled using non-specific membrane labeling, or "EVQuant_Template_Immuno.xlsx" for additional immune-specific labeling of EVs (latest versions will be available at https://github.com/MEvanRoyen/EVQuant and http://www.evquant.nl).
3. The template is used to store sample information and in combination with **Table 1** on page 5 can be used as a guideline for sample preparation.
4. Add each sample including internal standards (Conditioned medium and Liposomes) and negative controls (PBS and/or culture medium) to an Eppendorf tube (optional: samples can be added to a 96 deep well plate for preparing the samples using multichannel pipets).
5. **Non-specific membrane labeling only:** Prepare an BSA mix for all the samples by adding the calculated volumes of 0.3% wt/vol BSA (5 µl/sample) and PBS (5 µl/sample) to a 1.5 ml Eppendorf tube and add 10 ul of BSA mix to each sample.
6. **Non-specific and Immune-specific labeling:** Prepare an antibody mix for all the samples by adding the calculated volumes of immunofluorescent antibodies, 0.3% wt/vol BSA (be sure the final concentration of BSA is 0.03% in a 50 µl staining volume) and PBS to a 1.5 ml Eppendorf tube (Make use of Table 1 and the Excel template) and add 10 µl of antibody mix to the samples and incubate for at least 2 hours at RT in the dark.
7. Prepare a Rhodamine mix for all the samples by adding the calculated volumes of Rhodamine (2 µl/sample) and PBS (98 µl/sample) to a 50 ml falcon tube (always prepare 10-20% excess of mix).
8. EVs are non-specifically labeled by adding 100 µl of the Rhodamine mix to each sample (in case a deep well plate is used, this step can be carried out row by row using a multichannel pipet). Incubate the mixture for a minimum of 10 min at RT.
9. During the 10-minute staining procedure in step 8, the volume of Acrylamide/Bis solution for the TEMED mix (89.5 µl/sample) and APS mix (85 µl/sample) for the total number of samples can be transferred to two separate 50 ml falcon tubes and put at RT (always prepare 10-20% excess of mix).
10. When non-specific labeling is finished (step 8) add the total amount of TEMED and APS to the corresponding Acrylamide/Bis solution and gently swirl the tubes.
11. Add 90 ul of both TEMED mix and APS mix to the Eppendorf tube containing the EV mix and transfer 300 µl of the still aqueous solution in one of wells of the 96-wells glass bottom imaging plate (in case a deep well plate is used for sample preparation, this step can be carried out row by row using multichannel pipets).
12. Repeat step 11 for each sample.
13. After the last sample is transferred to the glass bottom 96-wells imaging plate, store the 96-wells plate at RT in the dark.
14. Image acquisition using a confocal microscope can be performed after 1h, but acquisition is even possible two days after preparation of the plate.

#### Table 1. EVQuant sample preparation

| Component | With antibodies (µl) | Without antibodies (µl) |
|---|---|---|
| | **N=1** | **N=1** |
| EV/Nanoparticle sample* | 1-40 | 1-40 |
| PBS | Adjust to 40 | Adjust to 40 |
| **Total sample** | **40** | **40** |
| CD63-alexa488 | 0.625 | 0 |
| CD9-alexa647 (0.2% BSA) | 2 | 0 |
| PBS | Adjust to 10 | 5 |
| BSA (0.3% w/v) | 3.7 | 5 |
| **Total antibody/BSA mix** | **10** | **10** |

| Component | With/Without antibodies (µl) |
|---|---|
| | **N=1** |
| **Sample + antibody/BSA mix** | 50 |
| PBS | 98 |
| Rhodamine (25 ng/ul) | 2 |
| **Total volume EV mix** | **150** |
| Acrylamide/Bis Solution 37,5:1 30% w/v | 89.5 |
| Ultra-pure Temed | 0.5 |
| **Total volume TEMED mix** | **90** |
| Acrylamide/Bis Solution 37,5:1 30% w/v | 85 |
| Ultra-pure Temed | **5** |
| **Total volume APS mix** | **90** |
| EV mix | 150 |
| TEMED mix | 90 |
| APS mix | 90 |
| **Total volume** | **330** |

*** In case of samples with a very low particle concentration, EV/Nanoparticle sample volume can be increased up to 140 ul. In this case the Rhodamine/Antibodies are directly added to the 140 ul of sample.**

#### 2. EVQuant: Setting up the microscope

When the EVQuant assay is performed for the first time, the microscope needs to be set-up which first includes defining a proper laser output and detection threshold to allow accurate EV quantification/characterization. A guideline to setup a laser scanning confocal microscope or a high-content screening system is described in part A. Because of its three-dimensional context, absolute quantification by the EVQuant assay also requires a one-time calibration of the imaging volume in which particles are detected for each microscope, as it depends on the unique optical specifications and settings of the (confocal) microscope that is used. A guideline to define the detection volume is described in part B. The calibration procedure described above needs to be carried out only once for each microscopic system and corresponding settings. If something is changed in the set-up of the microscope (e.g. due to maintenance) or the acquisition settings, the calibration procedure needs to be repeated.

##### (A) Defining the laser output and calibration of detection threshold

**1.** 30 minutes prior to sample preparation, turn on the microscope and lasers which will be used for the quantification/characterization

**2.** To find the right microscope set-up, the following samples are prepared and measured using different instrument settings;
   a. A minimum of three dilutions of fluorescent 100nm tetraspeck beads (Invitrogen) (e.g. 5, 2.5, 1.25 µl of beads)
   b. Negative control (dye only)

**3.** Prepare the calibration samples according to the EVQuant sample preparation protocol in a glass bottom 96-wells imaging plate (rhodamine dye is also added to the beads to mimic background noise)

**4.** Set-up the microscope for total EV quantification using the Rhodamine dye which has an excitation peak around 540 nm and an emission peak around 560 nm. The settings presented in **Table 2** can be used as a guideline to set-up a laser scanning confocal microscope or high content screening system.

**5.** Position the sample at the microscope and find the glass-sample interface manually (find the change from the dark signal to the brighter sample layer) or if possible, using an autofocus option

**6.** By stepwise changing the laser output (% of laser power/exposure time) above and below the pre-defined output (3-5 steps), a suitable laser output for accurate EV quantification can be identified during image analysis. Capture at least 10 images of each sample recorded at a height of 40 µm above the glass-sample interface for each laser output.

**7.** Analysis of the images is performed on a computer running the open source software Fiji (ImageJ) (https://imagej.net/Fiji/Downloads)

**Table 2: Microscope setups used for EVQuant on a laser scanning confocal microscope and a high-content screening system.**

| Parameter | Description: |
|---|---|
| Microscope: | Zeiss, LSM 510 Axiovert 200M inverted microscope |
| Running Software: | AIM version 3.2 |
| Objective: | 40x, NA 1.3, oil Plan-neofluar |
| Dichroic mirror 1: | HFT 488/543 |
| Dichroic mirror 2: | NFT 545 |
| Filter: | LP560 |
| Laser: | HeNe 543 |
| Laser output used: | 4.50 µW |
| Detector Gain: | 750 |
| Pinhole: | 1 airy unit |
| Pixel dwell time: | 1.60 µs |
| Average (Line): | 4 |
| Scan Zoom: | 5.0 |
| Image size: | 1024x1024 pixels (46.1 µm x 46.1 µm) |
| Pixel size: | 0.04 µm |

| Parameter | Description: |
|---|---|
| Microscope: | Opera Phenix, High-content screening system (Perkin Elmer) |
| Objective: | 40x, NA 1.1, water objective |
| Filter: | 570-630 |
| Laser: | 561 nm solid state laser |
| Laser output used: | 100% |
| Exposure time: | 500msec |
| Image size: | 2160x2160 pixels (323 µm x 323 µm) |
| Pixel size: | 0.150 µm |

8. Download the EVQuant plugin (latest version will be available at https://github.com/MEvanRoyen/EVQuant and http://www.evquant.nl). Open Fiji and install the plugin using the menus Plugin -> Install Plugin.
9. As the EVQuant plugin requires stacks of images, it is helpful to first create a stack of images for each sample by concatenating all the images corresponding to each same sample/laser output combination (Image>Stacks>Tools>Concatenate) and saving the stack as a new Tiff file.
10. Run the EVQuant threshold calibration plugin (Plugins>EVQuant>EVQuant Tools>EVQuant calibration (threshold)) and follow the instructions. The calibration procedure needs to be performed for one specific laser output at a time.
11. The plugin first quantifies the number of particles that are detected in the Negative Control (dye only) sample and step-wise increases the threshold used for selecting particles until the analysis has reached the point that the number of detected particles in the Negative Control sample is below the predefined threshold of 1 particle/2500 $um^2$.
12. After defining the threshold, the plugin quantifies the number of spots in the dilution samples and will show a summary and calibration curve of the analysis.
13. Save or copy the calibration results to an excel file. The calibrated threshold is also stored in the memory of Fiji and automatically filled in during subsequent EVQuant analysis procedures.
14. Repeat step 10 to 13 for the different laser outputs used
15. Select a laser output which shows a good linear relation between quantified particles and dilution.
16. From now on, this laser output and corresponding calibrated threshold will be used for subsequent EVQuant analysis.

#### (B) Calibrating the imaging volume

Due to the scattering of light, resulting in the typical point spread function (PSF), particles will be detected in multiple planes. To analyze the effect of the PSF on the effective optical slice thickness (EOST) in single plane measurements, multiple Z-stacks need to be acquired for 3D particle analysis in which both detected particles per plane and detected particle centers per plane are quantified. In this way, the effective optical slice thickness can be determined which allows a more accurate absolute particle quantification.

17. 30 minutes before sample preparation, turn on the microscope and lasers which will be used for the quantification/characterization
18. Load the microscope configuration which was chosen as the best set-up during the system calibration procedure
19. The detection volume is defined using;
    a. Fluorescent 100 nm tetraspeck beads
    b. Internal standard (EVs in condition medium)

20. Prepare the calibration samples according to the EVQuant sample preparation protocol in a glass bottom 96-wells imaging plate (rhodamine dye is also added to the beads to mimic background noise)
21. Position the sample at the microscope and find the glass-sample interface manually or if possible, using an autofocus option
22. For each sample, acquire a minimum of three z-stacks of at least 21 planes with an interval of 0.5 μm recorded at a height of 30-50 μm above the glass-sample interface having the center of the z-stack at 40 μm
23. Transfer the images to a computer on which the open source software Fiji (ImageJ) is installed.
24. Run the EVQuant EOST calibration plugin (Plugins>EVQuant>EVQuant Tools>EVQuant calibration (eost)) and follow the instructions.
25. The plugin first detects the number of spots in every plane of the z-stack using your input settings (use the detection threshold which was determined experimentally during the previous calibration step)
26. Next, the detected spots at different heights in the z-stack that belong to the same particle are connected and the z-coordinate of the particle centroid for each particle is determined.
27. For every z-stack, both the average number of detected particles and particle centers in each slice are used to calculate the effective optical slice thickness in which particles are detected using this microscope setup.
28. When multiple z-stacks have been analyzed an average effective optical slice thickness (EOST) is automatically calculated.
29. From now on, this EOST is used to calculate the absolute EV concentrations of the measured samples on this specific system

### 3. EVQuant: Total EV quantification using non-specific membrane labeling

1. 30 minutes before sample preparation, turn on the microscope and lasers which will be used for the quantification/characterization
2. Prepare all the samples including internal standards (cell conditioned medium and/or liposomes) and negative controls (PBS and/or serum free medium) according to the EVQuant sample preparation protocol (page 6) in a glass bottom 96-wells imaging plate
3. Position the sample at the microscope and find the glass-sample interface manually or if possible, using an autofocus option.
4. Load the microscope configuration which was previously set-up during the system calibration procedure (page 8).
5. Capture several images (at least 10) of each sample (including the internal standards and negative controls) at a height of 40 µm above the glass.

**Analysis of images using Fiji**

6. Transfer acquired images to the analysis PC
7. Open Fiji and open the EVQuant excel template which was used during sample preparation to obtain the information about sample volumes, dilution etc.
8. It is possible to analyze the images from all samples at once by opening all the images in Fiji and concatenating the images (keep track of the order of the samples) by choosing the following option in the taskbar; Image>Stacks>Tools>Concatenate.
9. Run the EVQuant analysis plugin (Plugins>EVQuant>EVQuant Analysis) and follow the instructions (it is possible to obtain the raw particle counts by selecting the “Show Raw Results” box).
10. In the second window, fill in the sample information. It is possible to select Negative Controls which means results will be corrected for the particles detected in the negative controls.
11. The total number of particles and absolute EV concentration is automatically calculated for each sample using the following formula given below.
12. Copy the Results (and/or raw data) to excel in the “ImageJ results” tab of the EVQuant template and fill in the microscope characteristics.

$$Particle\ concentration = N * \frac{V_0}{V_q} * \frac{V_{tot}}{V_s} * D$$

$N$ = Particles counted by Fiji
$V_0$ = Volume of 1ml in µm$^3$
$V_q$ = Total volume used for quantification (in µm$^3$)
$V_{tot}$ = Total volume used for sample preparation (in µl)
$V_s$ = Sample volume (in µl)
$D$ = Sample dilution

**Analysis of images acquired by the Opera Phenix microscope using the Harmony software**

6. Open the measurement in the Harmony Software (Perkin Elmer)
7. Run the EVQuant Analysis Sequence (details of the Analysis Sequence will be available at http://www.evquant.nl)
   a. Particles are detected using the “Find spots” function in the Harmony software to determine the local maxima in each image based on a set threshold. A region of interest (ROI) is drawn around each particle. The threshold is chosen in such a way that the number of detected particles in the samples is comparable to the number of detected particles when images were analyzed by the EVQuant plugin.
   b. The total number of detected particles and several other parameters (e.g. median relative spot intensity, image area) are calculated for each sample and can be exported to a text file.
8. Export the analysis file “PlateResults.txt” to an external hard drive and copy the results to excel in the “Harmony PlateResults” tab of the EVQuant template.
9. Provide the microscope/sample/acquisition characteristics in the EVQuant template
10. In cell W3 the average counts of the Negative Control samples can be defined to calculate corrected EV counts of all samples.
11. The absolute EV concentration is automatically calculated for each sample using the following formula:

$$Particle\ concentration = N * \frac{V_0}{V_q} * \frac{V_{tot}}{V_s} * D$$

$N$ = Particles counted by Fiji
$V_0$ = Volume of 1ml in μm$^3$
$V_q$ = Total volume used for quantification (in μm$^3$)
$V_{tot}$ = Total volume used for sample preparation (in μl)
$V_s$ = Sample volume (in μl)
$D$ = Sample dilution

### 4. EVQuant: Quantification of EV subpopulations by combining general membrane labeling and immune-specific labeling

1. Thirty minutes before sample preparation, turn on the microscope and lasers which will be used for the quantification/characterization
2. Prepare all the samples including internal standards (cell conditioned medium and/or liposomes) and negative controls (PBS and/or serum free medium) according to the EVQuant sample preparation protocol (page 6) in a glass bottom 96-wells imaging plate
3. Position the sample at the microscope and find the glass-sample interface manually or if possible, using an autofocus option.
4. Load the microscope configuration which was previously set-up during the system calibration procedure and change the set-up by adding the additional channels corresponding to the fluorophores which are used for the immune-specific labeling
5. As fluorescence differs between fluorophores and in addition is depended on antibody affinity and the number of epitopes of the protein of interest present on the EV membrane, laser output and antibody concentration should be defined experimentally using antibody dilutions and multiple laser outputs. The optimal settings of the LSM510 and Opera Phenix microscope used for the CD9 and CD63 immunofluorescent antibodies in this study are found in **Table 3** and can be used as a starting point.
6. Capture several images (preferably more than 10) of each sample (including the internal standards and negative controls) at a height of 40 µm above the glass

**Analysis of images using Fiji**

7. Transfer acquired images to the analysis PC
8. Open Fiji and open the EVQuant excel immuno template "EVQuant_Template_Immuno.xlsx" which was used during sample preparation to obtain the information about sample volumes, dilution etc.
9. The analysis requires a stack of images in which each dye/biomarker has a separate channel.
10. It is possible to analyze the images from all samples at once by opening the stacks of all samples in Fiji and concatenating the images (keep track of the order of the samples) by choosing the following option in the taskbar; Image>Stacks>Tools>Concatenate.
11. Run the EVQuant immuno analysis plugin (Plugins>EVQuant>EVQuant Immuno analysis) and follow the instructions (it is possible to obtain the raw intensity data and threshold calibration graphs by selecting the "Show Raw Results" box).

12. In the second window, fill in the sample information. It is possible to select Negative Controls which means the results will be corrected for the particles detected in the negative controls.
13. For all samples, the number of EVs representing each marker combination is quantified and if selected corrected for the number of particles having the same marker combination in the negative control samples. Fractions of specific subpopulations are also calculated by dividing the number of particles of the different subpopulations to the total number of detected particles. The analysis result of the liposome control sample shows the possible non-specific binding of antibodies and/or false positive detection rate.
14. Copy the Results (and/or raw data) to excel in the "ImageJ Results" tab of the EVQuant immuno template and fill in the microscope characteristics.

**Analysis of images acquired by the Opera Phenix microscope using the Harmony Software and RStudio**

9. Open the measurement in the Harmony Software (Perkin Elmer)
10. Run the EVQuant immuno Analysis Sequence (there is a 2 and 3-channel version of the macro) (details of the Analysis Sequence will be available at http://www.evquant.nl)
    a. Images are resized by cropping the boundaries of the image to reduce imaging artefacts such as fluorescent bleaching on the side of the images.
    b. Next, particles are detected using the "Find spots" function on the non-specific membrane labeling (ROI is drawn around each particle)
    c. For every particle, the mean fluorescent intensity of the ROI in each fluorescent channel (different biomarkers) is stored in the analysis file.
    d. A second ROI drawn around each particle, excluding the particle itself, is used to measure the fluorescent intensity of the surrounding of each detected particle in each channel which is also stored in the analysis file.
11. After running the Analysis Sequence, export the analysis file "objects_population - spots.txt" to an external hard drive.
12. Transfer the analysis file to a computer on which the open source software Rstudio is installed (https://www.rstudio.com/).
13. Open Rstudio and open the R Script for analysis of 2 or 3 fluorescent channels (e.g. "Immuno_analysis_3_channels_488_568_647_v1.R") which will be available at https://github.com/MEvanRoyen/EVQuant and http://www.evquant.nl. The R Script processes the intensity data from all individually detected spots for every sample in all channels at the same time and in the end provides a summary file of the analysis results.
14. Run the R Script line by line by first positioning the cursor on the first line of code (Ctrl+Enter to run the line of code on which the cursor is positioned). The comments in the R Script provide information about the steps which are

performed to identify and quantify the different EV subpopulations based on the immune-specific markers. The main steps of the script are explained here;

a. First, the working directory and location of the analysis file are defined.
b. For each channel, the individual spot intensities are corrected for the background intensities and are stored as relative spot intensities.
c. Subsequently, for each sample and separate biomarker channel, the values of spots that have a negative relative intensity are selected and combined with absolute values of the same negative relative intensities to create a gaussian distribution of background noise. Distribution of background noise for each sample and separate biomarker channel is plotted in a histogram and fitted using a Gaussian function.
d. Mean plus three times the standard deviation of the Gaussian fit is used as a threshold to define EVs being positive for each marker of interest.
e. Subsequently, for all samples, the number of EVs representing each marker combination is quantified and corrected for the number of particles having the same marker combination in the negative control sample. Fractions of specific subpopulations are calculated by dividing the number of particles of the different subpopulations to the total number of detected particles. The analysis result of the liposome control sample shows the possible non-specific binding of antibodies and/or false positive detection rate.
f. A summary file containing all sample data is exported to the working directory.

**15.** Open the EVQuant immuno template “EVQuant_Template_Immuno.xlsx” which was used during setup and import the summary data to the corresponding “RStudio results” tab (3 or 2 channel tab). Select the cell “D2” and select the following option in the taskbar; Data>from Text>Find and open the summary file and Click Next> select delimiter “comma”>Click Finish>Select existing worksheet.

**16.** Concentrations in the summary file are calculated based on an undiluted sample volume of 40 µl. If samples are diluted or other volumes were used, concentrations are automatically recalculated in column AK:AQ in the 3 channel tab or S:U in the 2 channel tab based on the values stored in column B and C.

**Table 4: Microscope setups used for Immuno EVQuant on a laser scanning confocal microscope and a high-content screening system.**

| Parameter | Description: |
|---|---|
| Microscope: | LSM 510 Axiovert 200M inverted microscope (Zeiss) |
| Objective: | 40x, NA 1.3, oil Plan-neofluar |
| Acquisition mode: | Serial channels |
| Dichroic mirror 1: | HFT UV/488/543/633 |
| Dichroic mirror 2: | NFT 545 |
| Filter: | LP560 |
| Laser channel 1: | HeNe 543 nm |
| Laser channel 2: | Argon 488 nm |
| Laser channel 3: | HeNe 633 nm |
| Emission filter channel 1: | LP 560 nm |
| Emission filter channel 2: | BP 505-530 nm |
| Emission filter channel 3: | LP 650 nm |
| Detector Gain (all): | 750 |
| Pinhole: | 1 airy unit |
| Pixel dwell time: | 1.60 μs |
| Average (Line): | 4 |
| Scan Zoom: | 5.0 |
| Image size: | 1024x1024 pixels (46.1 μm x 46.1 μm) |
| Pixel size: | 0.04 μm |

| Parameter | Description: |
|---|---|
| Microscope: | Opera Phenix, High-content screening system (Perkin Elmer) |
| Objective: | 40x, NA 1.1, water objective |
| Acquisition mode: | Serial channels |
| Laser channel 1: | 561 nm solid state laser |
| Laser channel 2: | 488 nm solid state laser |
| Laser channel 3: | 647 nm solid state laser |
| Emission filter channel 1: | BP 570-630 nm |
| Emission filter channel 2: | BP 500-550 nm |
| Emission filter channel 3: | BP 650-760 nm |
| Laser output channel 1: | 100% |
| Laser output channel 2: | 100% |
| Laser output channel 3: | 100% |
| Exposure time channel 1: | 500 msec |
| Exposure time channel 2: | 1500 msec |
| Exposure time channel 3: | 1000 msec |
| Image size: | 2160x2160 pixels (323 μm x 323 μm) |
| Pixel size: | 0.150 μm |

### 5. EVQuant: EV size characterization

The fluorescent intensity of individual particles depends on the number of fluorescent moieties incorporated into the membrane surface of the particle. As a result, the fluorescent intensity of a particle increases quadratically with the diameter which can be exploited for EV size analysis. However, a standard single plane EVQuant measurement results in the detection of particles present in different focal planes which hampers the sizing based on fluorescence intensity. Therefore, multiple Z-stacks need to be acquired in which the maximum intensity of the center of each detected particle is determined by fitting a Gaussian function in x, y and z through the measured intensities at different heights in the z-stack. A stepwise protocol for the size characterization is found below.

**1.** 30 minutes before sample preparation, turn on the microscope and lasers which will be used for the quantification/characterization.
**2.** Load the microscope configuration which was previously set-up during the system calibration procedure.
**3.** The size characterization is calibrated using liposomes with a known size (diameter) which are preferable characterized by (cryo-)EM and/or DLS.
**4.** Prepare all the samples according to the EVQuant protocol (page 6) in a glass bottom 96-wells imaging plate.
**5.** Position the sample at the microscope and find the glass-sample interface manually or if possible, using an autofocus option
**6.** For each sample, acquire a minimum of three z-stacks of at least 21 planes with a maximum interval of 0.5 µm having the center of the z-stack at a height of 40 µm above the glass-sample interface.
**7.** Transfer the images to a computer and open Fiji (ImageJ)

**Calibration of the sizing analysis using particles of a known size (diameter)**

**8.** Open the EVQuant size analysis template “EVQuant_Template_Size.xlsx” to store the results (latest version will be available at https://github.com/MEvanRoyen/EVQuant and http://www.evquant.nl).
**9.** For every size characterization experiment, the analysis needs to be calibrated using the images of a liposome/bead (calibration) sample with a known size.
**10.** Run the EVQuant sizing calibration plugin (Plugins>EVQuant>EVQuant Tools>EVQuant calibration (sizing) and follow the instructions (it is possible to obtain the raw intensity data and intensity histogram by selecting the “Show Raw Results”/”Show plot” boxes).
**11.** For each z-stack, 3D particle analysis determines the particle centers in each slice of the z-stack in the same way as during the calibration of the detection volume.
**12.** For size analysis, particles are selected which have their center in the zstack in the slices beginning from 1/4 of the total number of slices and ending in 1/4 of the total number of slices from the end of the zstack. This excludes the particles having the center of the particle close to the edge of the zstack.

**13.** For each particle the slice with the maximum mean intensity is determined. At this height of the z-stack, the pixel intensities in horizontal and vertical lines across the ROI are measured and fitted using a Gaussian function to define the coordinates of the pixel having the highest intensity in x and y. Next, the intensity of this specific pixel is measured at different heights in the z-stack and intensities are fitted using a Gaussian to determine the maximum intensity of the particle in Z.

**14.** To calculate the average maximum intensity in Z of all particles in the calibration sample, the intensities are plotted and fitted using a gaussian function.

**15.** Having the median maximum intensity of the particles in the calibration sample and as the average size of the calibration particles is known, a calibration factor (X) is automatically calculated which now allows converting fluorescent intensities to particle sizes (diameter in nm).

**16.** Save or copy the calibration results to the excel file in the “Calibration Sample” tab. The calibration factor is also stored in the memory of Fiji and automatically filled in during subsequent EVQuant size analysis procedures.

**EVQuant sizing analysis**

**17.** After calibrating the size analysis, size characterization can easily be performed for every sample.

**18.** It is wise to analyze multiple z-stacks per sample for more accurate size analysis.

**19.** Run the EVQuant size analysis plugin (Plugins>EVQuant>EVQuant Size analysis and follow the instructions.

**20.** First, the maximum intensity in the center of each particle is determined in the same way as described in the size calibration procedure.

**21.** Using the previously determined calibration factor, intensities are automatically converted to particle sizes and plotted in a histogram. When multiple z-stacks have been analyzed for the same sample, size-distributions are automatically combined which increases the number of particles in each bin.

**22.** Copy the results to the excel template in the “Sample X” tab.

### APPENDIX

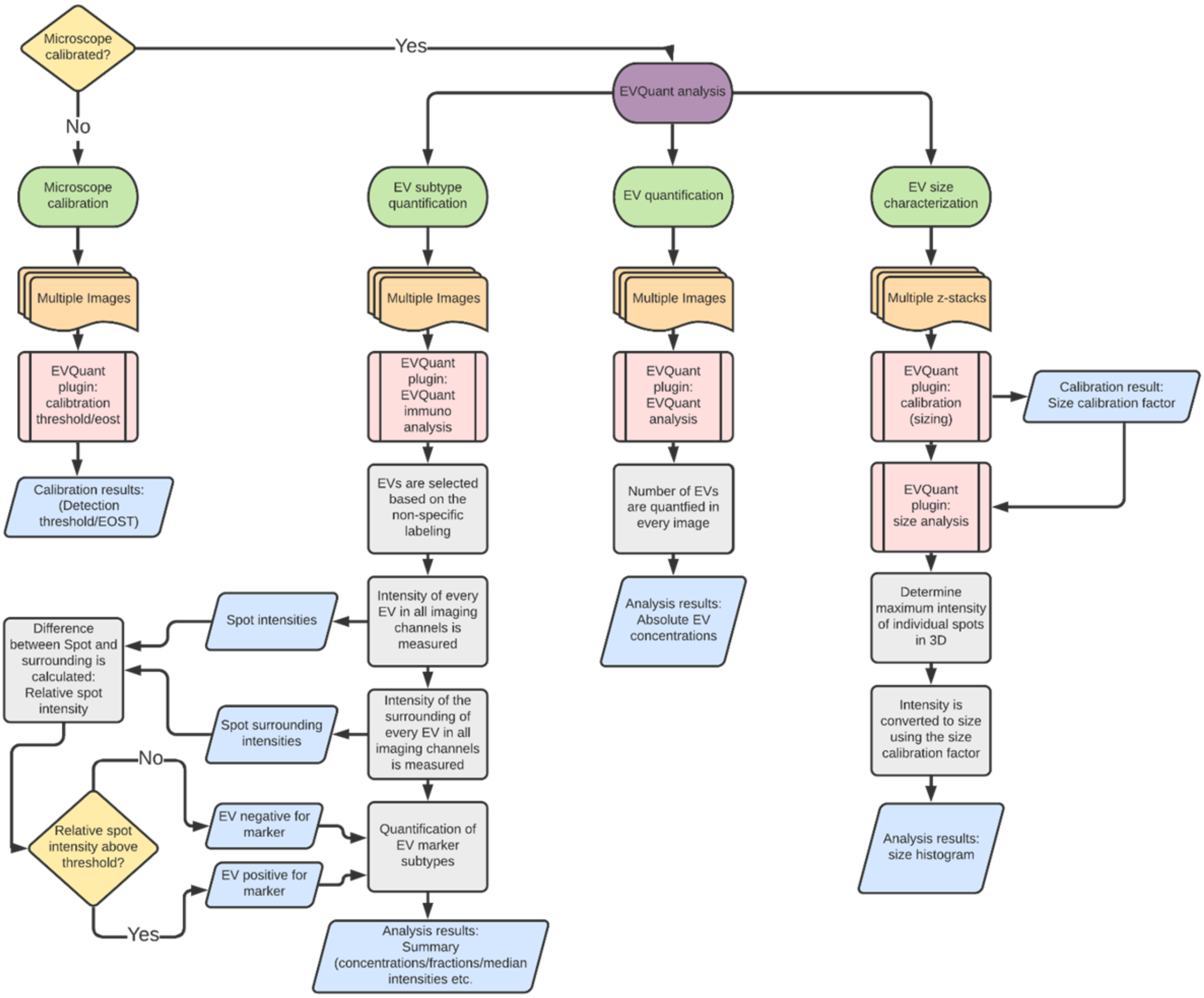


**Figure 1.** A flowchart presenting the different steps that are involved in the three different EVQuant analysis procedures.
